# Fast 3D imaging in the auditory cortex of awake mice reveals that astrocytes control neurovascular coupling responses at arteriole-capillary junctions

**DOI:** 10.1101/2024.06.28.601145

**Authors:** Barbara Lykke Lind, Andrea Volterra

## Abstract

Neurovascular coupling (NVC) increases blood flow, assuring adequate supply to active cortical regions by local redistribution via penetrating arterioles (PA) and branching capillaries. Astrocyte end-feet enwrapping these vascular structures possess machinery to regulate blood flow, but their participation in NVC is controversial. Via a new 3D+t two-photon imaging approach we visualized PA and capillaries simultaneously during naturally-occurring and tone-evoked dilations in the auditory cortex of awake mice. We observed that dilations occurred bidirectionally, and a fraction of them extended between compartments across the interconnecting sphincter, depending on the animal activity states. These multi-compartment dilations were preceded by rapid astrocyte end-foot Ca2+ signals around the sphincter. Reduction of this astrocytic Ca2+ activity in IP3R2KO mice suppressed multi-compartment dilations, revealing a pivotal role of pre-capillary sphincters in their bidirectional spread between vascular compartments under local control by astrocytes. This novel mechanism contributes to physiological regulation of laminar blood flow during NVC.

## Introduction

Adequate regulation of cerebral blood flow is the basis of healthy brain function. The neurovascular coupling (NVC) response ensures sufficient oxygen and energy during increased neuronal activity and, when impaired, contributes to pathophysiological alterations in several common cerebral diseases ^1^. The blood is recruited to the capillary bed from the brain surface via penetrating arterioles (PAs). The first branch point from the PAs into the capillary bed, the 1^st^ order capillary, also dubbed precapillary arteriole ^2,3^ or post-arteriolar transition zone ^4^, is believed to be critically important for blood flow regulation during NVC ^5^. Intriguingly, the presence of a gating structure, the pre-capillary sphincter, was recently described precisely at this level ^6^. This structure expresses α-smooth muscle actin in its mural cells (αSMA) ^6-8^, which makes it contractile and capable of modifying vessel diameter. Changes in the sphincter diameter may then control the blood flow to the downstream brain tissue ^9^. Astrocytes’ end-feet continuously cover all the blood vessel segments and could, in principle, impose control at each level ^10^. Intracellular Ca2+ increases in astrocytes can induce the release of vasoactive factors ^11^, yet the role of astrocytes in blood flow regulation remains controversial. Notably, it is unclear if Ca2+ elevations in astrocytes are rapid and prevalent enough to contribute to blood flow recruitment ^12^. To this point, different studies produced discrepant results: some reported that astrocytes recruit blood flow, but exclusively at the capillary level ^13,14^, others that astrocytes instead regulate PA diameter, but only during extended sensory experiences ^15^, and others that astrocytes do not exert any Ca2+-dependent regulation on blood flow. The latter conclusion was based on the authors’ inability to observe any changes in the NVC response in mice lacking IP3 receptor type-2 (IP3R2), the IP3R isoform thought to drive most of the intracellular Ca^2+^ elevations in astrocytes ^16-18^. Independent of their different conclusions, all the above studies looked at blood vascular dynamics in a simplistic manner, mainly focusing on individual compartments, and disregarding the heterogenous nature of dilations progressing through the vascular bed ^19-21^. Moreover, different studies used different imaging conditions to investigate astrocyte intracellular Ca^2+^ dynamics with respect to size, timing, and frequency of the recorded Ca2+ signals ^15,17,18,22-31^. Last but not least, most studies were performed in anesthetized mice in which both astrocytic Ca^2+^ ^32,33^ and blood flow responses ^34^ are severely dampened. Overall, these several shortcomings did not permit to draw firm conclusions on the role of astrocyte Ca2+ signaling in the control of brain hemodynamics.

In the present study, we aimed to overcome those pitfalls. To start, we performed our experiments in awake mice, which permitted us to consider the impact of the mouse activity state on the NVC response and the heterogeneity of the dilation patterns in different inherent conditions. Secondly, we used a 3D+t two-photon imaging approach ^32^, to our knowledge for the first time in NVC studies. With this approach, we could monitor entire regions (30-150 µm in the z-axis) at the branching point between penetrating arterioles and capillary bed and investigate simultaneously the vascular responses at the levels of PA, 1^st^ order capillary, and connecting sphincter, while also monitoring the Ca2+ activity in the surrounding astrocyte structures. Thirdly, we imaged these stacks at high speed (10 Hz), to capture the full range of astrocytic Ca2+ dynamics, including fast and local signals.

We report the existence of different patterns of natural dilations at the PA-capillary junctions in the mouse auditory cortex, which depends on the inherent activity of the mouse. Part of the observed dilations involved both PA and capillary, whereas others remained confined to only one compartment. Astrocytes occasionally underwent large Ca2+ activations, but they also often displayed smaller Ca2+ rises in the end-feet enwrapping the pre-capillary sphincters. Such local Ca2+ rises preceded spread of the dilations and specifically correlated with multicompartment dilations. Both the astrocyte Ca2+ activity and the spread of dilations across the sphincter were affected in mice lacking IP3R2 (IP3R2KO). When mice were subjected to auditory stimulation, dilation patterns were analogous to those seen in naturally behaving mice and similarly depended on astrocyte Ca2+ activity. We conclude that pre-capillary sphincters play a central role in the bi-directional spread of dilations between vascular compartments and that astrocyte Ca2+ signaling controls their function locally.

## Results

### 3D imaging of vessel dilations and astrocyte Ca2+ activity in the auditory cortex of awake mice: observations along the descending PA paths

We investigated the astrocytic contribution to hemodynamic responses in the auditory cortex of awake mice (Figure 1A) focusing at first on the PA tracts. We mapped the surface vessels to identify appropriate positions for imaging, where PAs dive into the tissue, i.e., where in the vascular tree, the freshly oxygenated blood enters the cortex (Figure 1B). By 2-photon imaging, we then studied astrocyte and blood vessel dynamics along the descending PA paths. Astrocyte Ca2+ activity was visualized using the *GFAPCreERT2:GCaMP6f* mouse strain that enables astrocyte-selective, inducible expression of the green Ca2+ indicator, GCaMP6f, upon tamoxifen (TAM) injections (Methods). In parallel, PA dynamics were revealed by loading blood vessels with red fluorescent dye (Texas Red-dextran) to highlight their lumen. In this first set of experiments, we positioned our three-dimensional fields of view (3D-FOV) along the PAs to cover descending tracts up to 280 µm in the z dimension (Figure 1C). Our fast 3D imaging approach ^32^ enabled us to monitor vascular and astrocyte dynamics quasi-simultaneously in multiple focal planes along the z-axis (Figure 1C, Sup. figure S1). Our investigation of the astrocyte dynamics focused on the Ca2+ activity within the end-feet, as they represent the points in which the astrocytes contact the contractile vascular cells. The end-feet regions of interest (ROIs) were automatically defined as the structures present within 2 µm from the vessel surface in each z-plane and repositioned during vascular dynamics, according to the vessel diameter changes (Methods and Sup. figure S1). In this way, we followed astrocyte end-foot Ca2+ activity along single PAs while tracing the volume changes occurring in the corresponding PA segments (Figure 1D). Some end-foot Ca2+ activities spread far along the PA and covered up to 205 µm in the z-dimension, whereas others stayed local and were seen just in a single focal plane (Figure 1E, Sup. figure S2A). As for the spontaneous PA dilations that we observed, only about half (324/668) were accompanied by Ca2+ activity in the end-feet along the PA. In these cases, the spread of PA dilations was larger than in the absence of the astrocyte Ca2+ activity and reached further down inside the cortex (Figure 1F). Moreover, it correlated with the spread of the end-foot Ca2+ activity (Sup. figure S2B). Since large astrocytic Ca^2+^ elevations are seen during mouse locomotion ^22,32,35^, we in parallel recorded EMGs with electrodes implanted in the mice’s neck muscles (Figure 1D, bottom). We found that the spread of the astrocytic end-foot Ca2+ activity in the z-direction almost doubled during mouse movements with respect to when the mouse was at rest, changing, on average, from 23±3µm to 50±7µm (Figure 1E).

**Figure 1.**
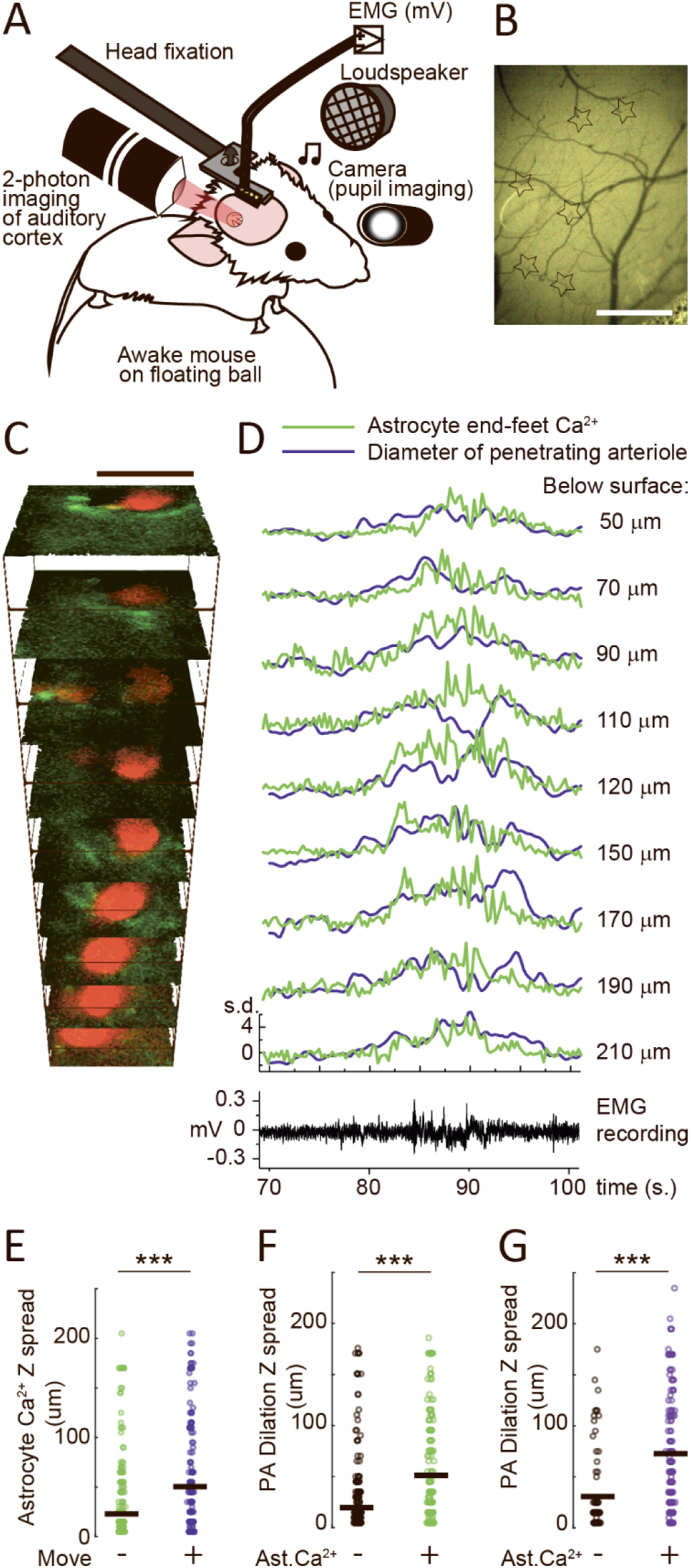
3D imaging of the PA dynamics in the auditory cortex of awake mice: both naturally occurring dilations and astrocytic Ca2+ activity spread along the descending PA path. A. Complete setup for our experiments in chronically implanted head-fixed awake mice. B. Brightfield image of the vessels at the surface of the auditory cortex seen from the chronic cranial window. Visual inspection of the blood flow direction in vessels allows distinction between arterioles and venules. *Stars*: PAs. Scale bar: 500 µm. C. Representative cumulative average of the 3D timeseries taken in the auditory cortex of a *GFAPCreERT2:GCaMP6f* mouse (green, reporting astrocyte Ca2+ activity) injected in the circulation with Texas red dextran (red, reporting vessels lumen). Imaging was performed in 20 µm z-steps, from 50 to 210 µm below the surface. Scale bar: 20 µm. D. *Top:* Example traces reporting simultaneous astrocyte end-foot Ca2+ activity (mean fluorescence) and PA diameter (area) changes normalized to SD values in the different z-planes of our 3D-FOV, before and during a period of mouse locomotion. The resting periods are used as baseline. *Bottom*: The movements are visible in the electromyogram (EMG) recorded from the mouse’s neck muscles throughout the imaging session. E. Spread of end-foot astrocyte Ca2+ activity along the length of the PA; *green*: mouse is resting; *violet*: mouse is moving (Mann-Whitney test: p=1.8x10^-11^; N=668 Ca2+ events in the resting mouse and 262 in the moving mouse). F. Spread of PA dilations along the length of the PA in the resting mouse; *black*: without Ca2+ activity in the astrocyte end-feet; *green*: with the astrocytic Ca2+ activity (Mann-Whitney test: p=5.7x10^-16^; N=324 dilations without astrocyte Ca2+ and 344 with Ca2+). G. Spread of PA dilations along the length of the PA as in F, but in the moving mouse; *black*: without Ca2+ activity in the astrocyte end-feet; *violet*: with the astrocytic Ca2+ activity (Mann-Whitney test: p=1.5x10^-9^; N=78 dilations without astrocyte Ca2+ and 184 with Ca2+). In E-G, data are from 8 FOVs: x=75-121 µm; y=53-111 µm; z=100-280 µm, in 10-20 µm steps from 6 mice.

In agreement with previous reports, we found that not all mouse movements triggered an astrocytic Ca2+ activity in the end-feet along the PA ^22^. Still, >70% of PA dilations seen during mouse movements (184/262) were associated to increased Ca2+ activity in the astrocyte end-feet, a higher proportion than the one seen in the resting mouse. As during rest, the PA dilations during movement that coincided with Ca2+ increases in the end-feet, also spread significantly further than those occurring solitary (Figure 1G). These data suggest that large end-foot Ca2+ activity influences PA dilations without being a necessary prerequisite for the dilations to occur.

### 3D imaging of vessel dilations and astrocyte end-foot Ca2+ activity in the auditory cortex of awake mice: observations at the PA-capillary junction

While PA dilations increase blood flow distribution to many cortical layers, a second and more specific level of regulation occurs at the capillary branching points at different cortical depths ^36^. An increased blood flow in one of the 1^st^ order capillary branches enhances the delivery to areas downstream of the branch ^5^. Recently, an investigation in the barrel cortex described a pre-capillary sphincter at the junction with the PA ^6^. Therefore, we decided to focus our study of NVC in the auditory cortex on this junction (Figure 2A). Via the luminal red fluorescence, we visualized the sphincter as a narrowing of the lumen proximal to the PA. In addition, in 50% of the junctions (n = 20), we noticed the presence of a bulb at the capillary edge, likely generated by a local reduction in the coverage by contractile cells ^37^. The shape of the PA-capillary junctions was variable, and some junctions had a less obvious indentation. Nonetheless, nearly all junctions showed some degree of pre-constriction and, irrespective of the shape, it is known that the entire 1^st^ order capillaries are contractile ^6,37^. Therefore, we decided to define all these connections between a PA and the 1^st^ order capillary as sphincters. Initially, we investigated the position of the astrocytic structures at the vascular junction using *GFAP-EGFP* mice with green-fluorescent astrocytes ^38^. The astrocytes were localized at the branching points (Figure 2A), covering most of the 1^st^ order capillary up to the sphincter and in contact with it (Figure 2B To detect the potential role of these astrocytes, we then performed dynamic studies in TAM-treated *GFAPCreERT2:GCaMP6f* mice, centering our 3D-FOV at the level of the PA-1^st^ order capillary junction. In these experiments, we selected 3D-FOVs much smaller than those utilized for studying dilations along the PAs and took focal planes at 1µm z-steps to ensure continuity in the local imaging around the vasculature (Sup. Figure S3). Thanks to fast two-photon scanning, we could investigate quasi-simultaneously the dilation patterns of the two vascular compartments and the rapid Ca^2+^ changes in astrocyte end-feet ROIs at both the arteriole and capillary, i.e. 1^st^ order capillary, levels. This setting enabled us to define location and temporal sequence of the astrocyte Ca2+ elevations relative to the vascular volume changes. In previous studies in awake mice, when recording from 3D-FOVs of dimensions similar to those used here, we had detected two different types of activities in astrocytic end-feet, involving either local or much larger Ca2+ events ^32^. The former were asynchronous and small (µm-scale), generally restrained within a single end-foot, whereas the latter expanded widely to occupy several end-feet in different z-planes. In the present recordings around the vascular junction, we similarly observed small and large events. Concerning the large Ca2+ elevations, they were rather infrequent, occurring every few minutes, and invaded ≥75% of the end-foot ROIs present in all z-planes of our 3D-FOV. We defined these events “large Ca2+ activities” (Figure 2C) and found that 71±7% of them overlapped in time with mouse movements (Sup. figure S2C). Since large astrocytic Ca^2+^ activities are known to occur during arousal states driven by noradrenaline release ^39^, and noradrenaline release is signaled by the animal’s pupil expansions ^40-43^, we monitored the mouse pupil diameter during the imaging sessions using an IR-camera. Thereby, we verified if large astrocyte Ca2+ activities coincided with pupil expansions. Indeed, while most pupil expansions occurred without a large astrocytic Ca^2+^ activity, when such an astrocyte activity took place, it almost always (92 ± 13% of the cases) coincided (within 1 sec) with a pupil expansion (Sup. figure S2D). However, as observed for the dilations along the PA path, the dilations around the junction were not triggered just by animal movement or by large astrocytic Ca2+ activities. Indeed, in the majority of cases (75.4±5.9%) dilations occurred when the mouse was at rest and without being accompanied by any large astrocytic Ca2+ activity. Looking at the temporal sequence of the events, when a large astrocytic Ca2+ activity was associated with a mouse movement, the Ca2+ rise in the astrocyte started after the onset of the movement (by 1.0 ± 1.25s, Sup. figure S2E and S2F). In some cases, it preceded the onset of the dilation, but, on average, it rose after the vessel dilation (by 2.3 ± 0.84s, Figure 2D). Therefore, this type of large end-foot astrocytic Ca2+ activity appears to be associated with movements and arousal states but not to be responsible for triggering the dilations seen at the PA-1^st^ order capillary junctions during the mouse movements.

**Figure 2.**
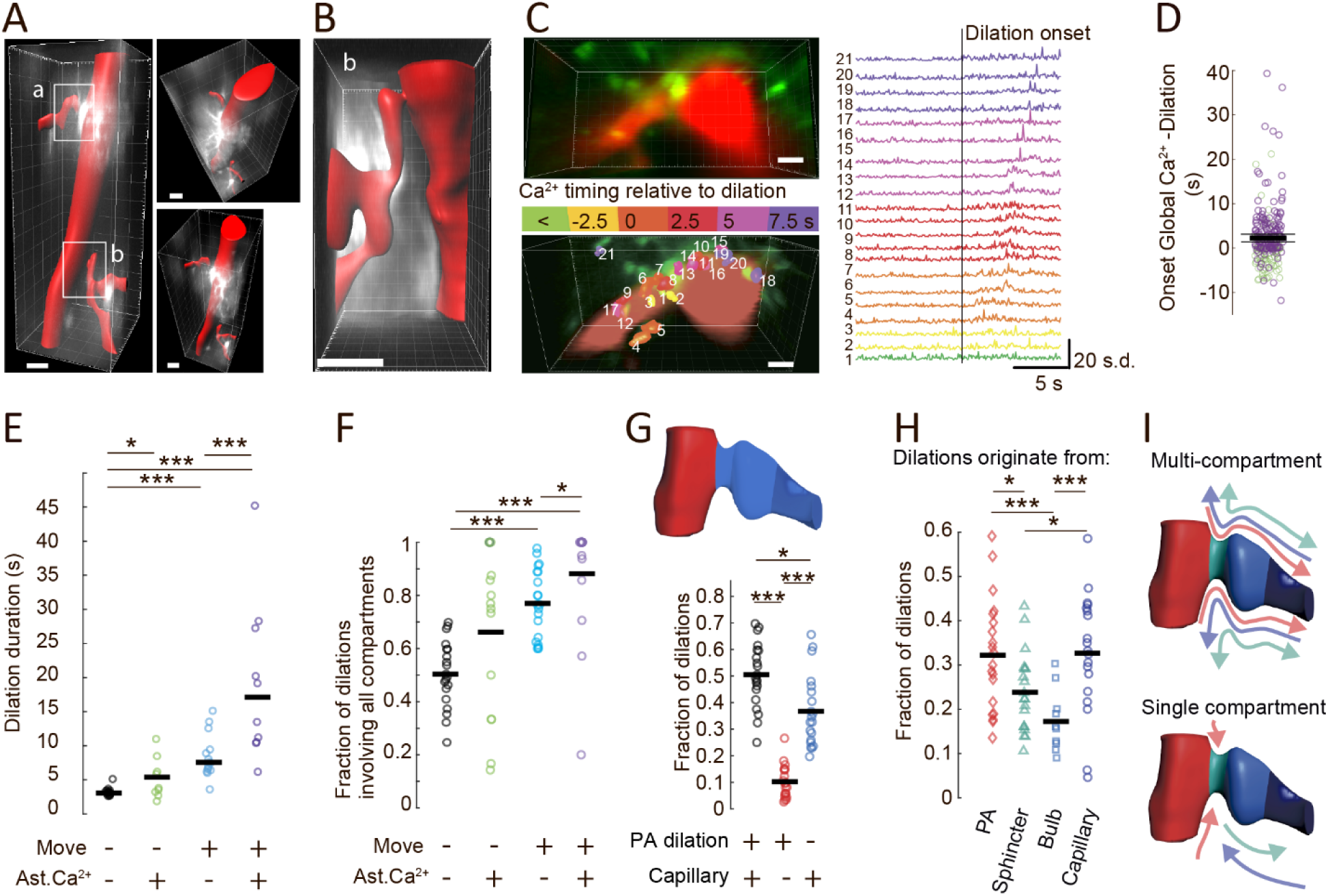
Spread of naturally occurring dilations in the auditory cortex of awake mice is gated at the sphincter between PA and capillary compartments. Correlation with end-foot astrocyte Ca2+ dynamics and animal motor state. A. Reconstruction of a 3D stack imaged live from 90 to 360 µm below the surface in the auditory cortex of an *EGFP-GFAP* mouse. Shown from the side (*left*) and from above (*right*). The reconstruction shows a penetrating arteriole delivering blood into the tissue and two 1^st^ order capillaries branching at different depths of the cortical layer II/III (a and b white squares on the *left*). *Red*: Texas red-dextran highlights vessels lumen; *white*: EGFP signal shows reconstruction of the astrocyte larger processes. Scale bar: 20 µm. B. Zoom-in on the 3D stack shown in A, depicting the lower branch (b) region (from 250 to 330 µm below the surface). Colors as in A. Scale bar: 20 µm. C. Representative cumulative average of a 3D time series, taken from a *GFAPCreERT2:GCaMP6f* mouse injected with Texas red-dextran in the vessel. The 3D-stack was imaged from 170 to 200 µm below the cortical surface, at 30 levels 1 µm apart. *Left, top*: cumulative signals show in red the vessel lumen and in green the neighboring astrocyte structures. *Left, bottom*: the same picture as above, but with multiple 3D-ROIs of astrocyte Ca2+ activity measuring a large astrocytic Ca2+ event associated with the animal’s locomotion and a dilation initiated at the pre-capillary sphincter. Ca2+ activity ROIs are color-coded to appreciate their temporal-spatial dynamics (Ca2+ onset time) relative to the dilation onset. The red signal is masked to show the 3D structure of the vessel. *Right*: Traces of mean astrocyte end-foot Ca2+ activity from each ROI. ROIs and traces are color-coded in relation to the timing of dilation onset. Scale bar: 8 µm. D. Delay between the onset time of all the large astrocytic end-foot Ca2+ activities and the onset time of the vessel dilations seen in the 3D-FOV when the mouse is resting (*green*) or is moving (*violet*). Mean Delay 2.3±0.86 s (Mean: *Thick black line,* 95% C.L. limits: *thin black lines*). N=223 dilation events in the moving mouse accompanied by large astrocyte Ca2+ activity. E. Average duration of dilations per 3D-FOV in different conditions. *Black*: in the resting mouse when no astrocyte Ca2+ activity is observed; 162±31.2 dilations/FOV); *green*: in the resting mouse in the presence of large astrocyte Ca2+ activity (4.2±2.0 dilations/FOV); *azure*: in the moving mouse when no astrocyte Ca2+ activity is observed (47.6±22.1 dilations/FOV); *violet*: in the moving mouse in the presence of large astrocyte Ca2+ activity (6.5±2.5 dilations/FOV). Dilations occurring in the resting animal were significantly longer when accompanied by an astrocyte Ca2+ event (t-test: without vs. with Ca2+, p=0.046), but significantly shorter than dilations occurring during mouse movements, accompanied or not by astrocyte Ca2+ (t-test: resting vs. moving without Ca2+: p= 6x10^-7^; resting vs moving with Ca2+: p = 3.3x10^-6^). Dilations during mouse movements were significantly longer when accompanied by an astrocyte large Ca2+ activity (t-test: moving vs moving with Ca2+: p= 0.0016). N=20 FOVs in 6 mice; all values were compared with ANOVA before t-test p=1.46x10^-9^. T-tests were paired, two-tailed, and p-values adjusted with a Holm correction for multiple comparisons. F. Average fraction of dilations per 3D-FOV comprising all the vessel compartments in the FOV in different conditions. Color coding, conditions, number of dilations analyzed/FOV, number of FOV, and number of mice are as in E. The fraction of dilations significantly increased when the mouse moved with respect to when it was at rest, both when a large astrocyte Ca2+ activity accompanied the movement and when it did not (Kruskal Wallis ANOVA (p=2.7 x 10^-6^) followed by Wilcoxon test with Holm correction: resting without vs. with Ca2+: p=0.059; resting vs moving, both without Ca2+: p= 0.00019; resting vs moving with Ca2+: p=0.00016). The fraction of multicompartmental dilations in the moving mouse further increased when movement was accompanied by an astrocytic Ca+ elevation (Wilcoxon test with Holm correction: moving without Ca2+ vs moving with Ca2+: p=0.013). G. Average fraction of dilations per 3D-FOV in the resting animal according to the involved vessel compartments. *Top*: Representation of the vascular compartments present in the 3D-FOV in all our experiments. The different vascular compartments have been subdivided and depicted in different colors: PA (*red*), capillary branch, comprising the sphincter, bulb and 1^st^ order capillary (*blue*). *Bottom*: Average fraction of dilations involving either all the compartments (multicompartmental, *black*) or each of them individually according to color coding. 162±31.2 dilations/FOV, N=20 FOVs, 6 mice. Half the dilations were multicompartmental, while the other half involved each compartment individually, with significantly more dilations occurring in the capillary branch (*blue*) compared to the PA (*red*). Friedmans ANOVA (p=1.59 x 10^-7^) followed by Wilcoxon test with Holm correction: multicompartment vs. PA: p =1.4 x 10^-4^; multicompartment vs. capillary branch: p= 0.015; capillary branch vs PA: p= 1.0 x 10^-4^. H. Average fraction of dilations per 3D-FOV in the resting mouse according to the compartment in which they originate: segmented compartments are depicted with different colors and symbols: PA (*red, diamond*), sphincter (*cyan, triangle*), bulb (*light blue, square*), capillary (*dark blue, circle*); 162±31.2 dilations/FOV, N=20 FOVs, 6 mice. Dilations had heterogeneous origin and direction, with significantly higher fractions originating from PA and capillary than from sphincter and bulb (Friedmans ANOVA (p=1.56x10^-7^) followed by Wilcoxon test with Holm correction: PA vs. sphincter: p = 0.032; PA vs bulb: p = 4.4 x 10^-4^; capillary vs sphincter: p = 0.025; capillary vs bulb: p = 1.3 x 10^-3^. I. Illustration of the direction of spreading of the dilations across the sphincter according to the compartment of origination (color-coding as in H). *Top*: multi-compartment dilations; *bottom*: single-compartment dilations. All data in C-I of this figure are from N=20 FOVs in 6 mice. 3D imaging coordinates: x=56-75 µm; y=15-44 µm; z=21-35 µm in 1µm steps.

### Bidirectional spread of dilations at the pre-capillary sphincter: dependence on the activity and arousal state of the awake mouse

During our repeated 3D-imaging sessions which were in most cases of 2 min duration, we detected many spontaneous vessel dilation events at the junction (Sup. Figure S3). Initially we thought that such events were part of the NVC response evoked by the natural sound perception, increasing excitation and metabolic needs of the auditory neurons. However, we noticed that the dilation events varied largely depending on whether they occurred when the mouse was moving or resting, as well as if a large astrocytic Ca^2+^ activity took place coincidentally or not (Figure 2E and F). Notably, dilations occurring when the mouse was moving lasted significantly longer (mean duration: 7.57 ± 1.32 s) than those in the resting mouse (3.06 ± 0.27 s, Figure 2E and S2G). Moreover, dilations in the moving mouse were in most cases (77%) multicompartmental, involving both the PA and the capillary branch (Figure 2F). In contrast, in the resting mouse, ∼50% of the dilations were mono-compartmental, confined to the vessel tract either upstream or downstream the sphincter. When dilations occurring in the moving mouse were accompanied by a large astrocytic Ca2+ activity, they lasted even longer (Figure 2E) and were almost always (92% of cases) multicompartmental (Figure 2F). Also in the resting mouse, coincidence of a large astrocyte Ca2+ event, increased the duration of dilations (Figure 2E), but without significantly increasing the proportion of multicompartmental dilations (Figure 2F). These data indicate that the animal movements, and to a lesser degree the arousal state, shape the hemodynamic responses in the auditory cortex, prolonging dilations at this junction in time and space.

Next, we analyzed the origin and direction of the dilations in the different conditions. Since only 51±5 % of the dilations during rest were multicompartmental, i.e., involved both PA and capillary, we evaluated which part of the vascular bed dilated alone in the remaining cases (Figure 2G). We found that monocompartmental dilations more frequently involved the capillary branch (37%±6% of all the dilations at rest), and only in 12±3% of cases the PA (Figure 2G). This heterogeneity was also reflected in the origin of the multicompartmental dilations: some of them first appeared at the PA, others at the capillary level and others at the junction. The latter ones were the only dilations initiating within our 3D-FOV; they represented about one third of all the multicompartmental dilations, and in most cases started at the sphincter, and just in a few cases at the bulb. The remaining dilations mostly originated outside the 3D-FOV; in 33 ± 5% of cases they were seen coming from the PA, and in 32 ±6% of cases from the capillary bed (Figure 2H). This heterogeneity in the sites of origin of the multicompartmental dilations did not depend on whether the mouse moved or was at rest, or the dilation was accompanied by a large astrocytic Ca2+ event or not. The site of origin did not influence the characteristics (volume and duration) of the dilations in each vascular compartment. In contrast, these were modulated by both the mouse’ behavioral state and the occurrence of large Ca2+ activity in astrocyte end-feet (Sup. figure S4). On the other hand, the fact that we observed both monocompartmental dilations stopping at the sphincter and multicompartmental ones crossing it in the resting mouse (Figure 2I), indicated to us that the pre-capillary sphincter is a site of control of the dilations’ spread. Such control was overcome during animal movements or arousal states, notably when these conditions were associated with large astrocytic Ca^2+^ activity, leading to the recruitment of both the upstream and the downstream vascular compartments, independent of the initiation site and the direction of the dilation. Based on these findings, we focused our next investigations on the astrocytic activity around the sphincter.

### Local Ca^2+^ activity in astrocytic end-feet at the sphincter precedes the arrival of multicompartment dilations

We considered that, when evaluating the role of astrocytes in the NVC response to auditory activation, the large astrocytic Ca2+ activities described above are likely confounding factors. According to recent data, they would be involved in associating brain states ^43^ or in integrating past events ^44^, functions that are not directly relevant to NVC initiation. Therefore, we decided to focus our next investigations on dilations and astrocyte Ca2+ elevations that are not associated to mouse movements/arousals, i.e. that occur during the animals’ resting periods (Figure 3). We hypothesized that astrocytes could exert their NVC regulation at the junction level and focused our attention on the astrocyte Ca2+ activity related to multicompartmental dilations spreading across the sphincter. By looking at time-averaged 3D images, we found heterogeneous local Ca2+ activities in the astrocyte end-feet and processes around the junction, which did not spread widely as for the large Ca2+ activities shown in Figure 2C. Following this initial observation, we decided to develop an automated analysis of the astrocytic end-feet Ca2+ activity in relation to the onset of spontaneous dilations. To establish the spatial-temporal sequence of these local astrocyte Ca2+ events with respect to the dilation events in an unbiased and accurate way, we aligned the 3D astrocyte Ca2+ imaging data to the onset of all the spontaneous dilations. We then identified voxels of interest (VOIs) with increased Ca^2+^ activity in the astrocytic end-feet (Methods and Sup. figure S5) and evaluated the probability that a Ca^2+^ increase occurred within these VOIs prior to a given dilation pattern (Sup. figure S5). Thereby, we identified VOIs that consistently responded in a defined temporal relation with a dilation (Figure 3A, Sup. figure S5), i.e., we identified the position and timing of pre-dilation Ca^2+^ activities in the astrocytic end-feet. Despite a certain variability in the VOIs positions and in the time-course of the related Ca2+ activities (Figure 3A), pre-dilation Ca2+ responses were consistently found primarily in the end-feet covering the sphincter (Figure 3B) and occurred at increased frequency when dilations were multicompartmental and spread across the sphincter (Sup. figure S6A). These Ca2+ responses did not depend on a specific origin and direction of the multicompartmental dilations (Figure 3B). Localized Ca2+ activity was also present in end-feet along the capillaries (Figure 3A and B). In contrast, we rarely saw pre-dilation Ca2+ activity in the end-feet on the PAs (Figure 3B). Regardless of their position, all the pre-dilation Ca^2+^ signals were local (9.7 ±3.8 µm z-axis) and short-lasting (0.98 ± 0.42 s) (Figure 3C). The interval by which pre-dilation astrocyte Ca2+ elevations at the sphincter preceded onset of dilations was longer when dilations arrived from upstream or downstream locations than when dilations started at the sphincter itself (Figure 3D and E). However, this longer anticipation might just depend on the fact that dilations coming from PA or capillaries initiated outside the FOV, i.e., earlier than observed. This been said, we observed that when a dilation that started at the capillary bed level and arrived at the sphincter was associated with Ca2+ events in both compartments, the astrocyte Ca2+ activity occurred first at the sphincter and then at the capillary (Figure 3D and F). This sequence did not visibly depend on spreading of the Ca2+ from the sphincter along the capillary (sup. Figure S6B). Rather, the two groups of end-foot astrocyte Ca2+ events appeared to be temporally separated and independent (Figure 3A, D-F and Sup. figure S6C), suggesting that Ca2+ activity at the sphincter is the primary pre-dilation event. Overall, the pre-dilation astrocyte Ca2+ activity at the sphincter showed anatomical and temporal characteristics in line with a possible role in controlling spreading of dilations across the sphincter.

**Figure 3.**
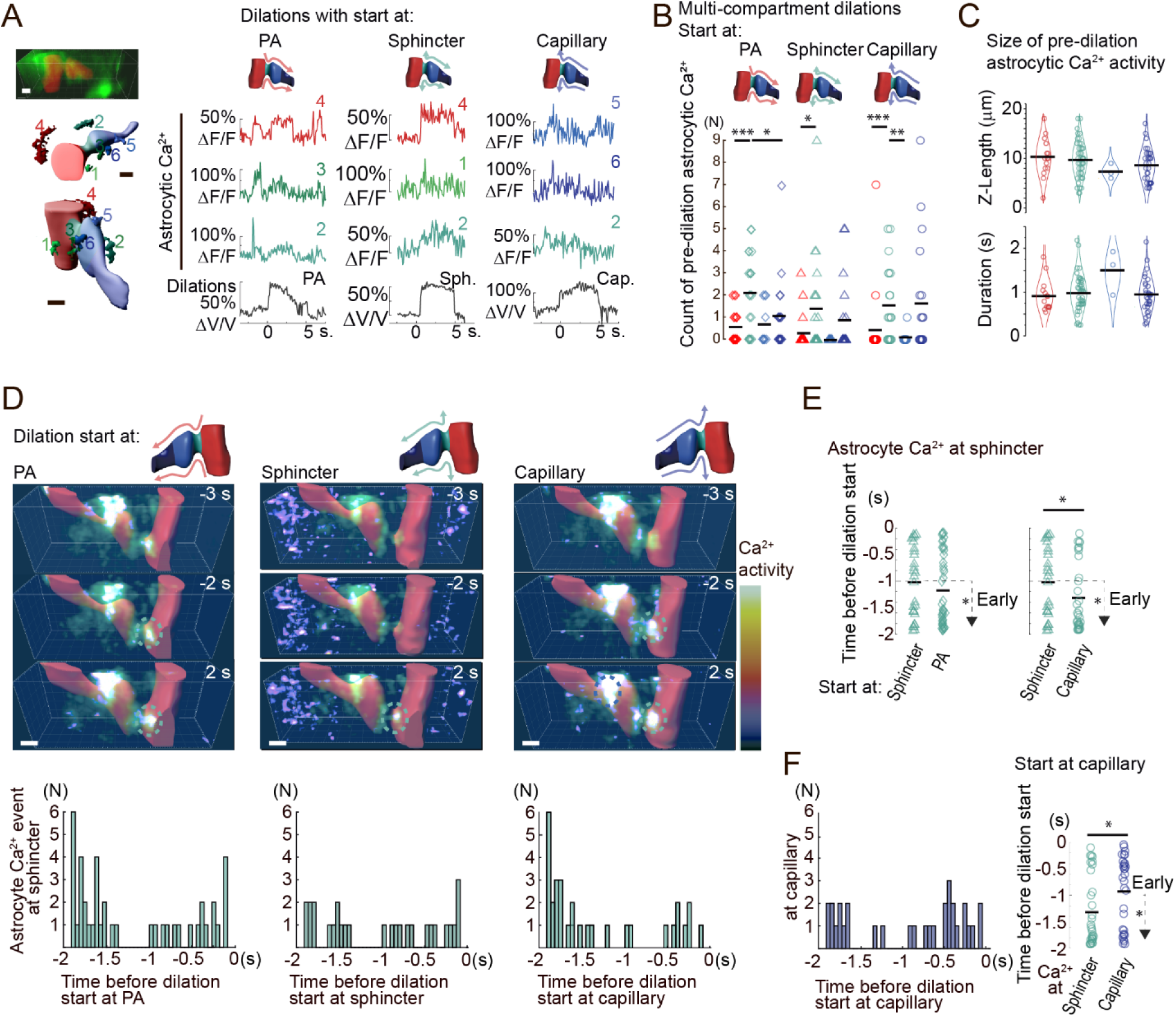
Local Ca^2+^ activity in astrocytic end-feet at the sphincter precedes multicompartment dilations arriving from distant vascular compartments in the resting mouse. A. Representative data showing local astrocyte end-foot Ca2+ activity preceding dilations originating in different compartments in our 3D-FOV (x=56.6 µm; y= 17.5 µm; z: from 194 to 216 µm below the brain surface in 1 µm steps). *Left, top*: cumulative average of the GCaMP6f (end-feet) and Texas-Red (vessel lumen) signals; *bottom*: the same vascular structures (*red*: PA; *cyan*: sphincter; *blue*: bulb/1^st^ order capillary) shown in two different orientations with several astrocytic VOIs at different locations around the junction, corresponding to the end-foot regions displaying pre-dilation Ca2+ activity shown in the traces on the *right*. Numbers identify each VOI; color identifies the VOI’s position on the vasculature. *Right*: Representative traces showing: *top*: time-course of astrocyte Ca2+ dynamics in VOIs numbered on the *left* during three dilations starting either at the PA (left column), the sphincter (middle column), or the capillary (right column); *bottom:* for each column, corresponding time-course of the dilation at the specific vessel segment where the dilation began. Pre-dilation Ca2+ activity appeared at several astrocyte end-foot locations, consisted of either individual peaks or more composite patterns, and was seen first in VOIs closest to the sphincter, including dilations arriving from the PA or the capillaries. Scale bars: 5 µm. B. Analysis of local pre-dilation end-foot astrocyte Ca2+ elevations associated with multicompartmental dilations. Quantification is by the number of end-foot VOIs with recurrent pre-dilation Ca2+ activity per FOV. Colors indicate the vascular location of the Ca2+ elevation (*red*: PA; *cyan*: sphincter; *light blue*: bulb; *blue*: capillary); symbols indicate the compartment of origin of the dilation (*diamond*: PA; *triangle*: pre-capillary sphincter; *empty circle*: capillary). Data are from 20 FOVs in 6 mice. End-foot Ca2+ events occurred more frequently at the sphincters, independent if the dilation initiated at the PA, the sphincter, or the capillaries. Tested with Kruskal Wallis ANOVA (KWA) followed by Wilcoxon test with Holm-correction for multiple comparisons: (a) dilation starting at PA: KWA p=0.019; count of end-foot Ca2+ events at sphincter vs. at PA p= 5.5x10-4; at sphincter vs. at bulb: p=0.062; at sphincter vs. at capillary: p = 0.027; (b) dilation starting at sphincter: KWA p=0.031; count of end-foot Ca2+ events at sphincter vs. at PA: p = 0.006; at sphincter vs. at bulb: p=0.25; at sphincter vs. at capillary: p = 0.41; (c) dilation starting at capillaries: KWA p=0.0033; count of end-foot Ca2+ events at sphincter vs at PA: p = 0.0011; at sphincter vs at bulb: p=0.048; at capillary vs at sphincter: p=0.21. The number of dilations/FOV starting at PA, sphincter and capillary, respectively, was 116.1±24.7, 82.4±19.7, and 67.2±16.2. N=20 FOV. C. Properties of the local astrocyte end-foot Ca2+ activities observed at four different vascular locations before multicompartmental dilations. Ca2+ activities are color-coded according to the vascular compartment where they are seen (*red*: PA; *cyan*: sphincter; *light blue*: bulb; *blue*: capillary). *Top*: spread of the end-foot Ca2+ activity along the z-axis (at PA: 10.1 ±1.9 µm; sphincter: 9.7 ±1.13 µm; bulb: 7.3 ±1.7 µm; capillary: 8.6 ±1.2 µm). *Bottom*: duration of the Ca2+ activity (at PA: 0.91 ±0.18 s; sphincter: 0.98 ±0.13 s; bulb: 1.5 ±0.58 s; capillary: 0.94 ±0.15 s). Data are from 20 FOVs in 6 mice. D. Timing of the astrocytic end-foot Ca2+ activity associated with multicompartmental dilations with different origin (*left*: PA; *middle*: sphincter; *right*: capillary bed). *Top*: Representative time-series of 3D-FOV (x= 75.5µm, y= 23.6µm taken from 95 to 125 µm below brain surface). Images are taken 3 sec before, 2 sec before and 2 sec after the dilation start. In *red*: Texas red-dextran masked signal highlighting vessel lumen; in *green*: cumulative astrocyte GCaMP6f signal highlighting active perivascular end-feet; *pseudocolor* scale: normalized Ca2+ activity levels to show Ca2+ changes per voxel. *Dotted line circles* highlight astrocyte Ca2+ activity at the sphincter (*cyan*) and at the capillary (*blue*). The activity is most precocious and intense when dilations arrive from PA or capillaries. Scale bar: 7 µm. *Bottom*: Distribution of the onset times of astrocytic Ca2+ signals seen at the sphincter before multicompartmental dilations starting at different vessel compartments. E. Comparison of the onset time of the different groups of pre-dilation end-foot Ca2+ activities shown in D. Following Kruskal Wallis ANOVA test we looked at individual comparisons. *Left*: timing of the pre-dilation Ca2+ signals for dilations starting at the sphincter vs at the PA. On average, there is no significant difference in onset time, but the number of early events (onset >1 sec before dilation) is significantly higher for dilations starting at PA (Mann-Whitney test; onset time: p=0.15; Wilcoxon test: count of early events: p=0.029). *Right*: same comparison but with dilations starting at the sphincter vs at the capillary. In this case, both onset time and number of early events are significantly different: end-foot Ca2+ activity starts earlier when dilations initiate at the capillary, and there are more early events (Mann-Whitney: onset time: p=0.027; Wilcoxon test: count of early events: p=0.043). The number of Ca2+ events detected before dilations starting at PA, sphincter and capillary, respectively, was N= 42, 28 and 31. The number of dilations/FOV analyzed starting at PA, sphincter, and capillary was 116.1±24.7, 82.4±19.7, and 67.2±16.2 respectively. All data are from 20 FOVs in 6 mice. F. Comparison of the onset times of astrocyte Ca2+ activities seen at the sphincter and at the capillary before arrival of multicompartmental dilations from the capillary bed. *Left*: Distribution of the Ca2+ signals at the capillary (*blue*). *Right*: The end-foot Ca2+ events around the sphincter started earlier and were more numerous in the early phases than those at the capillary (Onset time: Mann-Whitney test p=0.016 N=31 Ca2+ events; count of early Ca2+ events: Wilcoxon test, p=0.03, 67.2±16.2 dilations/FOV, from 20 FOVs in 6 mice.

### The astrocytic pre-dilation Ca^2+^ activity in end-feet at the sphincter is reduced in IP3R2KO mice

We then determined if the local pre-dilation astrocytic Ca^2+^ activity was IP3R2-dependent by repeating experiments in IP3R2KO mice that display reduced astrocyte end-foot Ca2+ activity in the *in vivo* awake condition ^45^. Indeed, in this group of mice, we observed a significant reduction in the number of local pre-dilation Ca2+ signals in astrocyte end-feet compared to wild-type mice. This reduction was seen in particular at the sphincter (Figure 4A-C, Sup. figure S7), while it was not significant at the PA and capillary (Sup. figure S7A and B). The reduction was most evident in association with multicompartmental dilations, independent of whether they arrived from the PA or the capillary (Sup. figure S7C), it involved particularly the earliest pre-dilation Ca^2+^ activity, namely the activity starting >1 s before dilation onset (Figure 4D; Sup. figure S7D) and occurring closest to the sphincter (Figure 4E; Sup. figure S7D). Overall, these results support the presence of an early, pre-dilation, IP3R2-dependent, Ca2+ increase in astrocytic end-feet, notably in the end-feet surrounding the pre-capillary sphincter, prior to the spread of dilations beyond the branch point.

**Figure 4.**
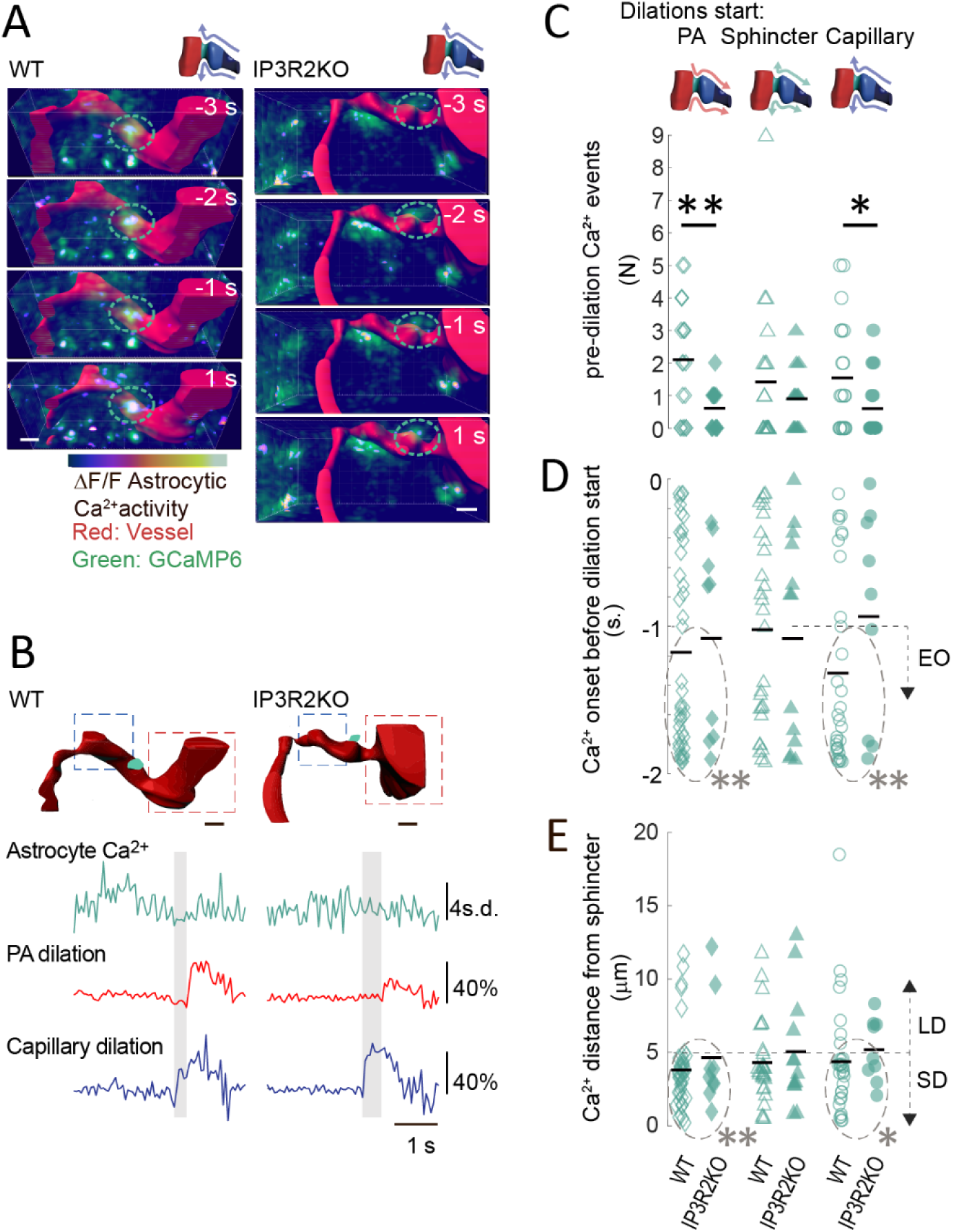
Pre-dilation astrocytic Ca^2+^ activity at the sphincter is suppressed in IP2R2KO mice; the most affected events are the earliest ones and those closest to the sphincter. A. Pre-dilation astrocyte end-foot local Ca2+ dynamics in wild-type (WT) and IP3R2KO mice. To illustrate the recurrence of the pre-dilation Ca2+ signals, their averaged activity in association with 3 spontaneous dilations is shown in representative 3D-FOVs in the period from 3 sec before to 1 sec after the start of the vasodilations. All vasodilations start at the capillary bed and spread to the PA. Each image is the cumulative average of 1 s interval in a 3D-FOV in: WT (*left*, x=75.5 µm, y=23.6 µm, z=170-200 µm below the cortical surface), or IP3R2KO (*right*, x=75.5 µm, y=30.7 µm, z=138-162 µm below the surface). In *red*: masked vessel lumen; in *green*: cumulative astrocyte GCaMP6f showing active end-feet; *pseudocolor scale*: normalized Ca2+ activity. *Dotted line circles* highlight astrocyte Ca2+ activity at the sphincter. Scale bars: 7 µm. B. Comparison of pre-dilation astrocyte Ca2+ signals at the pre-capillary sphincter before multicompartment vasodilations in WT and IP3R2KO mice. *Top*: astrocyte Ca2+ signals and vasodilations are measured in two ROIs spatially related to the vascular segments undergoing vasodilation: in the case shown, the vasodilation starts at the 1^st^ order capillary level (*blue*) and spreads to the PA (*red*). *Bottom*: representative astrocyte Ca2+ and vessel dilation traces in the defined ROIs. In the IP3R2KO mouse, the pre-dilation Ca2+ signals are reduced, and the PA dilation is delayed. C. Number of astrocyte Ca2+ events seen at the pre-capillary sphincter before multicompartmental dilations in several experiments in WT (*open symbols*) and IP3R2KO (*solid symbols*) mice. *Black lines* are mean values. Separate comparisons are shown for dilations starting at PA (*diamonds*), sphincter (*triangles*) or capillaries (*circles*). The number of pre-dilation Ca2+ events was significantly reduced in IP3R2KO vs. WT mice for dilations starting at PA or capillary, but not for dilations starting at the sphincter (Mann-Whitney test: WT vs KO: (a) dilations with origin at PA: p = 0.0077; (b) dilations with origin at sphincter: p = 0.45; (c) dilations with origin at capillary: p=0.029. Number of dilations/FOV analyzed with origin at PA, sphincter or capillary was, respectively: in WT: 116.1±24.7, 82.4±19.7 and 67.2±16.2 in N=20 FOVs in 6 mice. In IP3R2KO: 130.3±35.4, 76.9±21.9, and 50.9±14.3 in 14 FOVs in 4 mice. D. Timing of the pre-dilation astrocyte Ca2+ events seen at the sphincter in the 2 sec preceding multicompartment dilations in WT and IP3R2KO mice (*open* and *solid symbols*, respectively). Evaluation of the difference in onset time is conducted on the entire population of events and followed up by quantification at the level of the sub-population of earlier-onset events (EO, starting >1 before dilations start). Comparisons are performed separately for dilations starting at PA, sphincter or capillaries (*symbols* as in C). Despite no significant difference emerged between WT and IP3R2KO in the onset time of the entire population, the number of EO events was significantly reduced in IP3R2KO mice (*dashed grey circles*) for dilations starting at PA or capillary (Mann-Whitney test, WT vs KO: Dilations starting at (a) PA: onset time: all events: p=0.71; EO events: p = 0.0049; (b) at sphincter: onset time: all events: p= 0.79; EO events: p = 0.18; (c) at capillary: onset time: all events: p=0.13; EO events: p = 0.0075). The number of dilations/FOV analyzed with origin at PA, sphincter and capillary were, respectively: in WT: 116.1±24.7, 82.4±19.7, and 67.2±16.2 in 20 FOVs in 6 mice and in IP3R2KO: 130.3±35.4, 76.9±21.9 and 50.9±14.3 in N=14 FOVs in 4 mice. E. Distance from the sphincter of the pre-dilation astrocyte Ca2+ events in the 2 sec preceding multicompartmental dilations in WT and IP3R2KO mice (*open* and *solid symbols*, respectively). Comparisons are performed for dilations starting at PA, sphincter or capillaries (*symbols* as in C). *Black lines* are mean values: no statistical difference emerges between WT and IP3R2KO in any group of dilations. However, this could depend on the wide spatial dispersion of the events. Thus, for further analysis, events were sub-divided in two groups: those occurring at shorter distance (SD) from the sphincter (<5 µm) and those occurring at longer distances (LD, 5-20 µm). SD events were significantly reduced (*dashed grey circles*) for dilations starting at PA or capillary (Mann-Whitney test: WT vs KO: for dilations starting at (a) PA: SD: p=0.0038; LD: p=0.18; (b) sphincter: SD: p=0.52, LD p=0.18; (c) capillary: SD: p=0.023, LD p=0.15). The number of dilations with origin at PA, sphincter and capillary were, respectively: in WT: 116.1±24.7, 82.4±19.7 and 67.2±16.2 in 20 FOVs in 6 mice; in IP3R2KO: 130.3±35.4, 76.9±21.9 and 50.9±14.3 in 14 FOVs in 4 mice.

### Spread of dilations across the sphincter is impeded in IP3R2KO mice

We next assessed whether the reduced pre-dilation astrocyte Ca2+ activity at the sphincter in IP3R2KO mice had an impact on the vascular dynamics around the sphincter. In WT mice, we had observed that the pre-dilation Ca2+ activity was associated in particular to multicompartmental dilations, independent if they arrived from the PA or the capillary bed. Supporting a role for the astrocyte activity in controlling those dilation events specifically, we counted a decreased total number of multicompartmental dilations (-7%) and a proportionally increased number of single compartment dilations (+10%) in IP3R2KO compared to WT mice (Sup. figure S8A). In addition, we found that the proportion of multicompartmental dilations arriving at the junction from outside the 3D-FOV (either from the PA or the capillary compartment) was reduced in IP3R2KO mice, while the proportion of those initiated within the 3D-FOV (at either the sphincter or bulb) was increased (Figure 5A). Notably, it was the number of monocompartment dilations arriving from the capillary branch which increased (Sup. figure S8A), suggesting that less of them could spread across the sphincter to the PA in IP3R2KO mice (Figure 5B). We then discovered a second significant effect of IP3R2KO when assessing the speed by which the residual multicompartmental dilations spread from one compartment to the next one (Figure 5C). In WT mice, dilations spread from PA to capillary, on average, in 488 ± 18 ms, and, in the opposite direction, from capillary to PA, in 667 ± 17 ms (Figure 5C). Thus, the speed of compartmental transition was significantly higher when a dilation progressed from PA to capillary (608 ± 57 µm/s) than in the opposite direction (112 ± 5 µm/s, Sup. figure S8B). In IP3R2KO mice, dilations arriving from the PA took on average 260 ms longer in engaging the capillary than in WT mice. Likewise, dilations arriving from the capillaries, were delayed by 140 ms in crossing the sphincter and reaching the PA (Figure 5C, Sup. figure S8B). In the case of dilations initiated at the pre-capillary sphincter, spreading to the capillary was also delayed (by 110 ms) in IP3R2KO compared to WT mice (Figure 5C, Sup. figure S8B). In all cases, the slower spread of dilations was not due to local structural changes occurring in IP3R2KO mice, such as an increased distance separating PA and capillary structures compared to WT mice (Sup. figure S8C). These disturbances in the vascular dynamics of IP3R2KO mice had not been described before. In fact, several publications reported no changes in blood flow regulation in IP3R2KO with respect to WT mice ^16-18^. Interestingly, if we performed our analysis without taking into account intercompartmental spread and directionality of the observed dilations, but simply adding all the dilations together, we also obtained that average frequency (Sup. figure S9A) was not modified in IP3R2KO compared to WT mice. While all the above measures were done in the resting mouse, we observed additional effects produced by IP3R2KO during spontaneous movements of the animals. During movements we counted fewer dilation events (Sup. figure S9A) as well as a reduced intercompartmental spread of the dilations compared to WT mice (Sup. figure S9B-C). Thus, the astrocyte Ca2+ regulation of vascular dynamics appears to operate both during resting periods and movements of the mice. Overall, in IP3R2KO mice, both the pre-dilation Ca^2+^ activity in astrocytic end-feet surrounding the sphincter and the speed and efficiency of spread of the dilations across the sphincter were reduced. The latter phenomenon is likely responsible for the failed recruitment of additional vessel compartments and the reduced number of multicompartmental dilations, with a qualitative alteration in blood redistribution.

**Figure 5.**
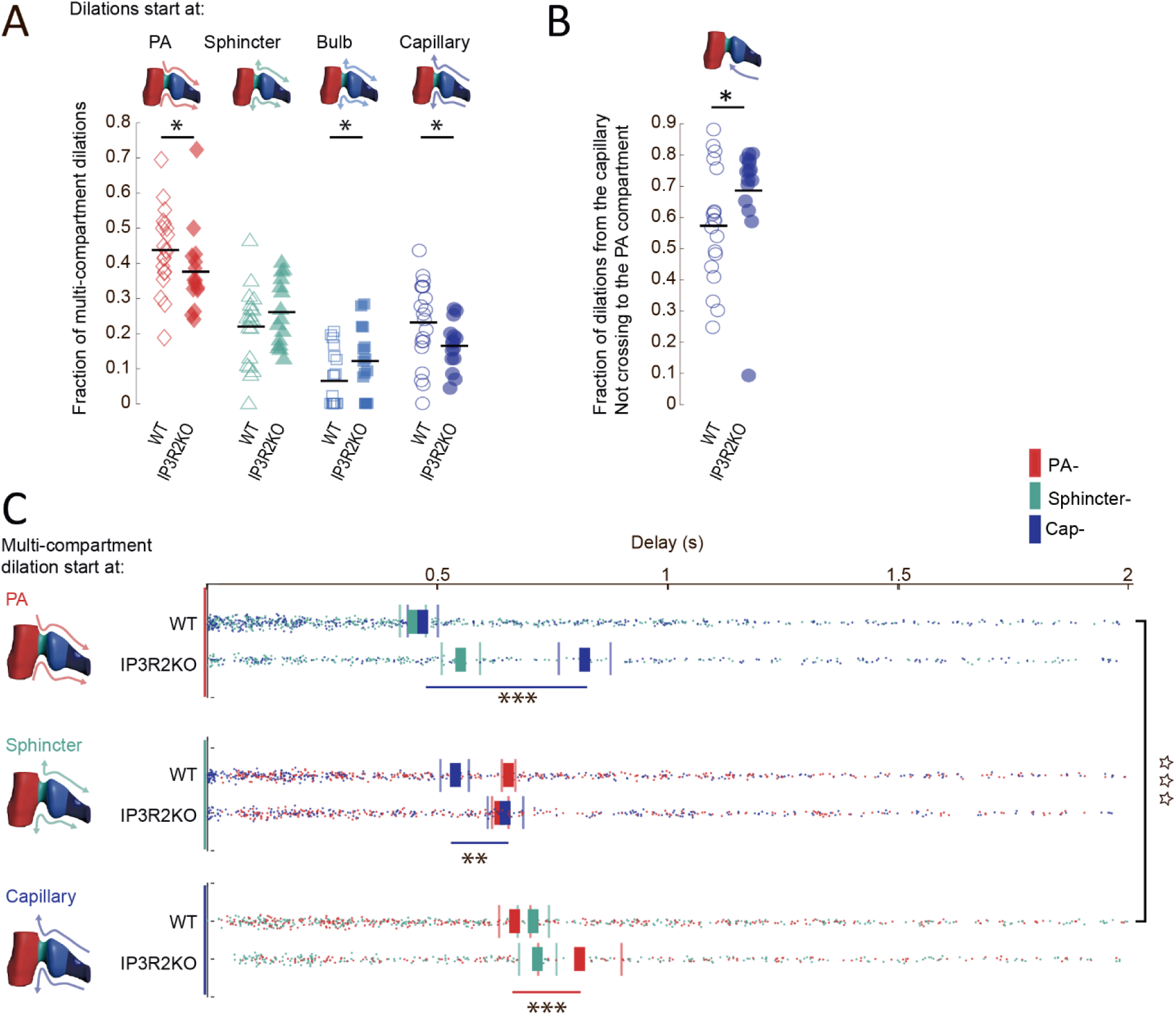
The spread of dilations from one vascular compartment to another is impaired in IP3R2KO mice. A. Fraction of multicompartment dilations according to their initiation site in WT and IP3R2KO mice. WT: *open symbols*; IP3R2KO: *solid symbols*. Dilations starting at PA: *red diamonds*; at sphincter: *cyan triangles*; at bulb, *light blue squares*; at capillaries, *blue circles*. *Black lines* are mean values. In IP3R2KO mice, the fraction of multicompartment dilations starting at the PA or the capillary bed was reduced compared to WT mice (First level statistical analysis: Friedmans ANOVA: WT: p=2.9x10^-7^, X^2^=33.2; KO: p=0.00022, X^2^=19.5; X^2^ test indicates more equal distribution of dilation origin in KO mice. Second level analysis: Mann-Whitney test: WT vs KO: dilations at PA: p = 0.017; at sphincter: p = 0.16; at bulb: p = 0.044; at capillary: p= 0.025). The number of dilations/FOV analyzed in WT was 82.65±20.13 in N=20 FOVs in 6 mice; in IP3R2KO, 75.27 ±16.73 in N=14 FOVs in 4 mice. B. Characteristics of the dilations arriving from the capillary compartment in WT and IP3R2KO mice. WT: *blue open circles*; IP3R2KO: *blue solid circles*. *Black lines* are mean values. In IP3R2KO mice, the fraction of dilations that did not cross the sphincter to become multicompartmental was larger than in WT mice (Mann-Whitney test: WT vs IP3R2KO: p = 0.027). Number of dilations/FOV analyzed: in WT: 166.84±30.41 in N=20 FOVs in 6 mice; in IP3R2KO: 176.2±41.24 in N=14 FOVs in 4 mice. C. Delay in the spread of multicompartmental dilations between vessel segments in WT and IP3R2KO mice. Dilations are presented according to their initiation segment (*top*: PA, in *red*; *middle*: sphincter, in *cyan*; *bottom*: capillary, in *blue*). Color-coded dots along the x-axis mark the start time of dilation of a given vessel compartment, which is measured as delay from the start time of the initiating compartment. The color-coded rectangles are the corresponding mean values, and the surrounding thin lines show the SEM. In WT mice (20 FOVs from 6 mice), dilations spread faster from PA to capillary than in the opposite direction, from capillary to PA (*Open stars* in vertical; Wilcoxon signed-rank test, p = 4.4 x 10^-6^; n = 366 dilations spreading from PA to capillary and 323 dilations from capillary to PA). In IP3R2KO mice, the spread of dilations across the sphincter is slower than in WT mice for dilations arriving from outside the FOV, regardless of the initiating compartment. Also, dilations starting at the sphincter are slower than in WT mice in reaching the capillary (Mann-Whitney test; WT vs. KO: (*top*) initiation at PA: spread: to sphincter: p=0.62; to capillary: p = 0.0000048; n = 366 dilations in WT and 142 in IP3R2KO mice; (*middle*) initiation at sphincter: spread: to capillary: p = 0.0017; to PA: p = 0.62; n = 329 dilations in WT and 239 in IP3R2KO mice; (*bottom*) initiation at capillary: spread: to PA: p = 0.0083; to sphincter: p = 0.93; n= 323 dilations in WT and 145 in IP3R2KO mice. Data from 20 FOVs in 6 WT mice and from 14 FOVs in 4 IP3R2KO mice.

### Auditory stimulations induce dilations and local astrocytic Ca2+ activity at the sphincter resembling those occurring naturally in resting mice

Our investigations so far focused on naturally-occurring dilations at the pre-capillary sphincter in the resting animal, excluding contributions by movement/arousal-induced mechanisms. However, these investigations did not directly demonstrate that the observed vascular responses belong to NVC, nor excluded that they depend on factors generated by sound perception other than the local neuronal activation ^46^. Thus, next we induced vascular responses via direct auditory stimulation and compared their properties to those of the “natural” dilations. The auditory cortex is formed by zones that gradually change sensitivity to different pitches ^47^. Therefore, we tested several tones and used intrinsic optical signaling (IOS) detection to determine the tonalities responsible for stimulating the cortical region exposed in our craniotomy. With IOS, we could also establish rough tonality maps and identify where, in a given cortical area, tone frequencies activated maximum blood flow increase (Figure 6A). To relate tone stimulations (10 sec) and associated IOS-blood flow responses (IOS maps) to local dilations at PA-capillary junctions (Figure 6A-B), we used combinations of three tones as stimulus, predicting that they would trigger a response in at least one of the cortical regions present in our cranial window. Initially, we investigated the correlation between wide-field IOS maps and individual vessel regulations at a locus in the auditory cortex containing four PA-capillary junctions. We sampled in parallel the individual vessel responses at the four branch points (one in layer I and three in layer II/III) present on three PAs entering the tissue at different cortical locations. The corresponding dilations differed in origin and directionality, but all had duration in line with the strength of the IOS signal detected in the same area: longer-lasting dilations were associated with tones producing strong local IOS responses, and shorter-lasting ones with tones producing weak IOS responses (Figure 6B). Noteworthy, the layer I junction showed less tone specificity than the layer II/III junctions, possibly because the flow pattern in superficial layers is less restrictive than in the deeper layers downstream. We found fewer junctions in the loci that we selected for the next observations. To incorporate all of them in our study, we tried a classification based on the individual IOS maps, i.e., we tried to link the observed dilation pattern at a given PA-capillary junction to the strength of the related tone-evoked IOS response. In particular, we considered the position in which the PA entered in the cortex with respect to the IOS map, i.e., whether its path and the related PA-capillary junction were at the center of the IOS map or displaced from it (Figure 6C). We found that most of the tones induced IOS responses somewhere in our cranial window, but not all these responses were centered on the region where a PA entered the cortex. When a tone induced a strong IOS response in the area of a junction (Figure 6C left), most of the dilations were long-lasting and quickly progressed across the pre-capillary sphincter to become multicompartmental, independent if they began at the capillary or the PA. In cases when the junction area was not at the center of the tone-evoked IOS response, we could still observe dilations in our 3D-FOV, but they mainly arrived from the capillaries (rarely from the PA) or originated at the sphincter (Figure 6C right). Tone-induced dilations varied largely in duration, ranging from >60 seconds to <1 sec (Sup. figure S10A). If the junction area was located at the center of the IOS response, dilations were at least 5 s-long and progressed beyond the sphincter to an extent proportional to their duration. The shortest ones were primarily restricted to the capillary bed, while the longest ones always propagated to all compartments (Sup. figure S10B), resembling the “natural” dilations in the resting animal (Figure 2). Also, the proportion of tone-evoked dilations that started at each location (respectively upstream of the sphincter, downstream, or at the sphincter) was analogous to that of the “natural” dilations (Sup. figure S10C). These observations strengthen the case that naturally occurring dilations in the resting animal directly reflect the auditory experience and represent the associated NVC response.

**Figure 6.**
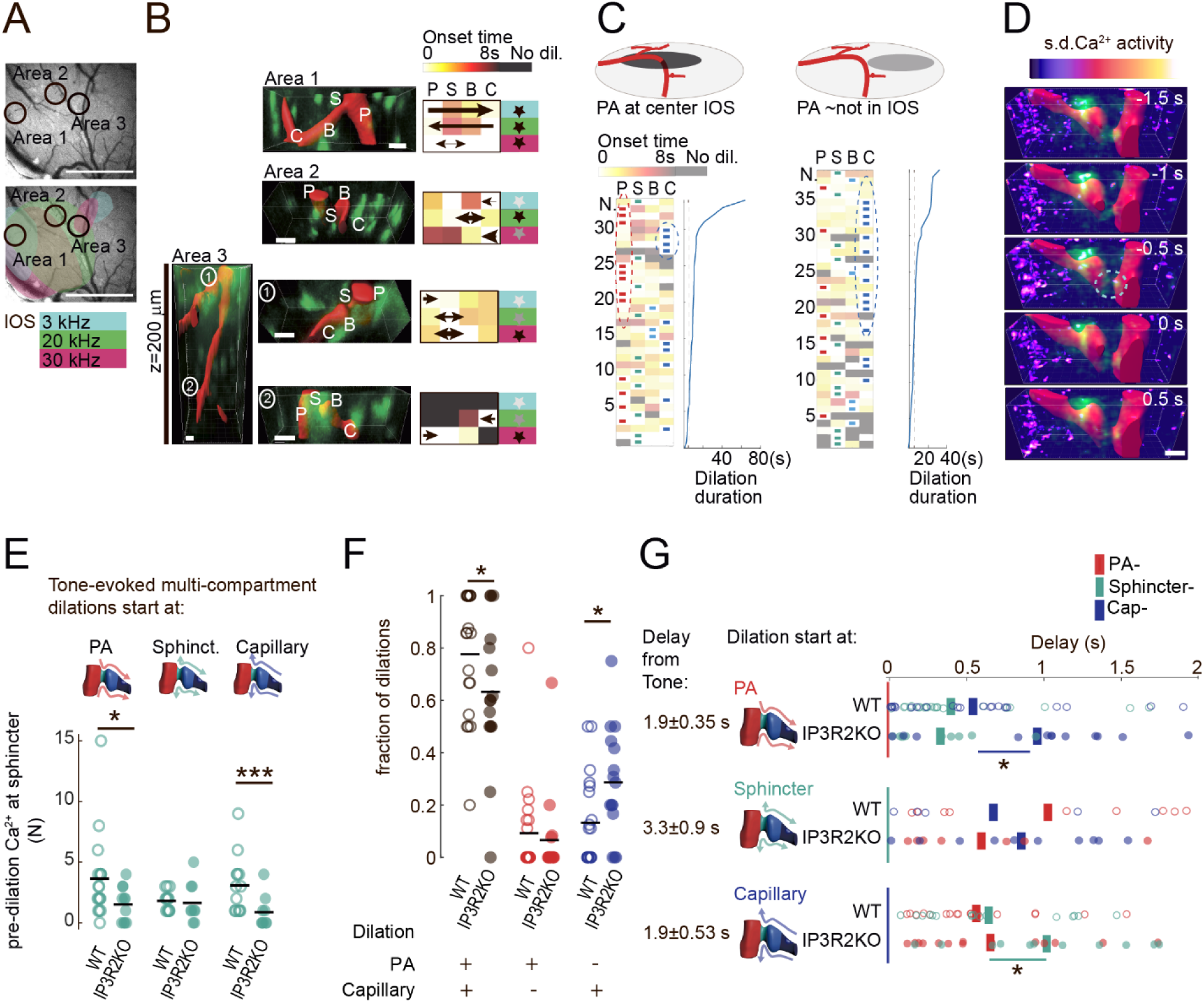
Pre-dilation astrocytic Ca^2+^ activity at the sphincter and vascular dilation patterns after auditory stimulations resemble those naturally occurring in resting mice and, like them, are altered in IP3R2KO mice. A. *Top*: Brightfield image of the brain surface in a chronic craniotomy over the auditory cortex. *Bottom*: Intrinsic optical signal (IOS)-based map of the tonality in the same auditory cortex region, with an overlay of three colors, each one marking the response pattern to the stimulation of the corresponding tonality (*azure*: 3kHz; *green*: 20 kHz; *red*: 30 kHz). The map shows that the three tone frequencies activate distinct but partly overlapping regions. Black circles identify the areas where three PAs enter the tissue (see panel B), indicating the tone-specificity of the blood flow response in each area. Scale bar: 1mm. B. Representative dilations at PA-1^st^ order capillary junctions in response to tone stimulations. Recordings are from 4 3D-FOVs in the 3 auditory cortex areas shown with circles in panel A at the center of which is the entrance point of the PAs. *Left*: images show vascular and astrocytic structures in the 4 3D-FOVs in the 3 areas and are cumulative averages of 2-min-long 3D timeseries; in *red*: Texas red-dextran masked signal highlighting vessel lumen; in *green*: astrocyte GCaMP6f signal highlighting the active perivascular end-feet. The 3D-FOVs in Areas 1 and 2 contain a single pre-capillary junction each, with the branching of the capillary from the PA at the level of layer II/III (170-200 µm and 177-202 µm below the cortical surface, respectively). The 3D-FOV in Area 3 contains 2 junctions (circled 1 and 2, respectively): the upper one in layer I (1, 64-86 µm below surface), and the lower one in layer II/III (2, 194-216 µm below surface). The different vascular sections are marked as: P for PA; S for sphincter; B for bulb; and C for capillary. *Right*: schematic representation of dilations evoked by tones of different frequencies (tones color-coded as in panel A). The timing of individual vessel segment dilation is depicted on a white-to-dark-red temporal scale (black indicates no dilation). The direction and duration of the dilations are schematized by the direction and thickness of the arrows, respectively (longer duration = thicker arrow). The position of the PA with respect to the IOS response evoked by a given tone frequency stimulation is summarized by color-coded stars: *black*, PA at the center of the IOS response; *grey*, PA at the border of the IOS response; *light grey*, PA further away from the IOS border. The length of the dilations correlated with the tone frequency that elicited the strongest IOS response in the specific 3D-FOV. Vessel dilation durations: Area 1: 3kHz stimulation: 40.7s; 20kHz: 15.4s; 30kHz: 3s. Area 2: 3kHz: 3.5s; 20kHz: 25.5s; 30kHz: 7.2s. Area 3, upper junction 1: 3kHz: 13.5s; 20kHz: 6.5s; 30kHz: 63.8s; lower junction 2: 3kHz: no dilation; 20kHz: 12.4s; 30kHz: 10.3s. The junction in layer I showed less tone specificity than the three junctions in layer II/III (see color-coded map in A). Scale bars: 10µm. C. Dilation patterns in the average responses to different tones in all the 3D-FOVs included in this study (n=73 tone responses in 20 FOVs in 6 mice). *Left:* patterns associated with tones that evoked a strong IOS response around the PA; *right:* patterns associated with tones that evoked weak or no IOS response around the PA. Dilation responses are ordered according to their duration (*blue plots* on the right of each column), from bottom (shortest) to top (longest). Timing of dilation of individual vessel compartments is represented in white-to-dark-red temporal scale as in panel B. The different vascular sections are marked as: P for PA; S for sphincter; B for bulb; and C for capillary. The vessel segment where the dilation originated is marked by a rectangular symbol in the following color code, PA: *red*; sphincter: *cyan*; bulb: *light blue*; capillary: *blue*. Dilations with the longest duration were evoked by tone stimulations with IOS response centered around the PA (*left*). They began at either the capillary (circled by *blue dotted line*) or PA (circled by *red dotted line*) and quickly progressed across the sphincter. In contrast, short dilations were evoked in areas with approximately no IOS response and often did not involve all compartments (*right*). Those involving all compartments mostly originated from the capillary (circled by *blue dotted line*). D. Representative time-series of pre-dilation astrocytic Ca2+ activity associated with tone-evoked multicompartmental dilations. The 3D-FOV, x= 75.5µm, y= 23.6µm, goes in z from 194 to 216 µm below the brain surface. Images are from 1.5 seconds before the spread of a multicompartmental dilation starting at the PA to 0.5 seconds after its start. *Red*: Texas red-dextran masked signal highlights vessel lumen; *green*: cumulative astrocyte GCaMP6f signal highlights active perivascular end-feet. *Pseudocolor*: Normalized astrocytic Ca2+ activity. *Dotted line circle* shows astrocytic Ca2+ activity at the sphincter prior to dilation onset. Scale bar: 10µm. E. Number of pre-dilation astrocyte end-foot Ca2+ activities seen at the sphincter before tone-evoked multicompartment dilations in WT and IP3R2KO. WT: *cyan, open circles*; IP3R2KO: *cyan, solid circles*. *Black lines* are mean values. Comparisons are presented in separate columns according to the site of origination of the dilation. The number of pre-dilation Ca2+ signals was significantly reduced in IP3R2KO compared to WT mice both for dilations that began upstream (PA) and downstream (capillary) the sphincter (Mann-Whitney test: WT vs. KO: dilation starting at PA: p = 0.026; at capillary: p=0.0046) but not for those initiated at the sphincter (WT vs KO: p = 0.22). Number of dilations/FOV starting at PA, sphincter, and capillary, respectively, were in WT: 1.98±0.62, 0.85±0.38, and 1.25±0.58, from N=20 FOVs in 6 mice; in IP3R2KO: 1.4±0.68, 1.3±0.83, and 1.13±0.63 from N=14 FOVs in 4 mice. F. Fraction of multicompartmental and single compartment dilations evoked by auditory stimulations in WT and IP3R2KO mice. WT: *open circles*; IP3R2KO: *solid circles*. *Black lines* are mean values. Analysis is restricted to dilations lasting >5 sec. Comparisons are presented for the ensemble of multicompartmental dilations (*black circles*), and separately for single-compartment ones involving the PA (*red circles*), or the capillary (*blue circles*). Multicompartmental dilations were significantly reduced in IP3R2KO mice, whereas single compartment dilations confined to the capillary were increased (First level statistical analysis: Friedmans ANOVA: WT: p=4.7x10^-6^, X^2^=24.5; KO: p=0.00032, X^2^=16.3. Second level analysis: Mann-Whitney test: WT vs KO: multicompartmental dilations: p=0.049; single compartment involving: the PA: p=0.52; the capillary: p = 0.012). The number of dilations/FOV analyzed in WT was 7.55±1.79 in N=20 FOVs in 6 mice; in IP3R2KO: 9.67±2.52 in N=14 FOVs in 4 mice. G. Temporal progression of tone-induced multicompartmental dilations arriving from outside the FOV in WT and IP3R2KO mice. WT: *open circles*; IP3R2KO: *solid circles*. Representation as detailed in Fig. 5C, according to the starting compartment: *red*, PA; *cyan*, sphincter; *blue*, capillary. Data are normalized to the arrival of the dilation in the FOV, but the average start point after the tone stimulation for each compartment is reported on the y-axis. The spread of the dilations to and across the sphincter was delayed in IP3R2KO mice (Kruskal Wallis followed by Mann-Whitney test; WT vs KO: (*top*) for dilations starting at PA: onset time at sphincter p = 0.55; at capillary p =0.019. N= 43 dilations in WT and 21 dilations in IP3R2KO; (*middle*) for dilations starting at sphincter: onset time at PA: p=0.82; at capillary p=0.37. N= 17 dilations in WT and 20 dilations in IP3R2KO; (*bottom*) for dilations starting at capillary: onset time at sphincter: p = 0.032; at PA: p = 0.19. N= 29 dilations in WT and 17 dilations in IP3R2KO. Data from 20 FOVs in 6 WT mice and form 14 FOVs in 4 IP3R2KO mice.

As a next step, we investigated whether tone-evoked dilations, like “natural” dilations, were preceded by local astrocyte end-foot Ca^2+^ activity. For this analysis, we used the same unbiased automated approach used for extracting the Ca2+ dynamics preceding “natural” dilations (Figure 3). In this case, our inclusion criterion was less permissive due to the much lower number of tone evoked dilations compared to the “natural” ones (Sup. figure S5, Methods). Nonetheless, we could consistently identify Ca2+ signals occurring mainly at the pre-capillary sphincter, preceding multicompartmental dilations (Figure 6D, Sup. figure S10D and S10E).

### The tone-evoked astrocyte Ca2+ responses and the blood vessel dilations are affected in IP3R2KO mice

To directly address the impact of the local astrocyte Ca2+ activity on the progression of tone-evoked dilations, we repeated the experiment in IP3R2KO mice. In WT mice, the large majority (79%) of the dilations arriving from either the PA or the capillary compartments were preceded by an astrocytic Ca2+ activity at the sphincter. This proportion decreased to 48% in IP3R2KO mice and was reflected in fewer pre-dilation events (Figure 6E). Similar to what was observed for the spontaneous dilations, in IP3R2KO mice the fraction of tone-evoked dilations that could not cross the sphincter and remained confined to the capillary bed was more than twice that in WT mice (28.7 ± 10.8% vs. 13.2 ± 7.4% in WT, Figure 6F). In complement, we observed that the progression of tone-evoked dilations across the sphincter in IP3R2KO mice was slowed down. Dilations arriving from the PA were delayed by 416 ± 145ms in reaching the capillary. Dilations from the capillary bed were also delayed by 376 ±208ms, mostly in recruiting the sphincter (Figure 6G). We conclude that IP3R2 deletion has a negative impact on tone-evoked dilations similar to the one seen in naturally occurring dilations. These data indicate that astrocytes, by their local Ca2+ activity at the sphincter, act as physiological regulators of the spreading of dilations between vascular compartments as part of the NVC response to auditory stimulation.

## Discussion

In this study, we identified a new role for astrocytes in NVC as regulators of the spread of dilations between the PA and capillary compartments in response to sound perception in the auditory cortex. The astrocyte control is exerted focally, via end-foot Ca2+ elevations occurring at the pre-capillary sphincters that connect the two compartments. Such local astrocyte activity precedes the arrival of dilations at the sphincters and controls their contractility, determining how much and how far blood is distributed during cortical activation. We obtained this information in the awake mouse thanks to fast volumetric two-photon imaging of vascular and astrocyte Ca2+ dynamics in large FOVs comprising the PA and capillary compartments and the connecting junction. By centering our 3D-FOVs on the vascular tree and the associated astrocytic end-feet and peri-vascular regions, we could capture even the most transient local Ca2+ activities and their interplay with the vascular dilations throughout the imaged vascular tracts. Only a minority of preceding studies were performed in the awake mouse, and none of them used fast volumetric imaging. This new approach enabled us to study all the natural dilation patterns occurring in a behaving mouse, and also to establish their origin, directionality, and level of progression between imaged compartments, revealing a previously unappreciated complexity. Thanks to this methodological advance, we could, on the one hand, overcome several of the shortcomings that affected previous work and alimented multi-year controversies on the role of astrocytes in NVC (see Introduction), and on the other, revisit the few recent studies performed in the awake mouse ^15,17,22,31,48^, complementing their findings and providing a more comprehensive picture. Among others, we could here dissociate the components of the natural astrocyte and vascular responses in the auditory cortex evoked by sound perception from those depending on the activity state of the mouse during locomotion and arousal. Given the large analogy between natural responses seen in the resting animals and responses evoked by natural tone stimulations, we conclude that the natural responses at rest are directly induced by the sensory experience and most likely represent the physiological NVC response to auditory activation.

The dilation patterns observed in our study differ significantly from those reported in previous work in anesthetized mice, characterized by quite stereotyped hemodynamic responses ^7,49^ and less correlation with the neural activity ^50^. Here, in our naturally behaving mice, the dilation patterns were highly heterogeneous and incorporated contextual contributions from the animal’s brain state, or inputs from other cortical areas, such as the motor cortex. During our recordings, mice moved and underwent arousals, and these activities were associated with large astrocytic Ca2+ events not seen in anesthetized mice ^32^. Such large astrocytic Ca2+ elevations were reminiscent of those recently shown to prolong the duration of PA dilations during sustained whisker stimulation in the barrel cortex of awake mice ^15^. Since during arousals, more dilations spread across the pre-capillary sphincters than in the resting mouse, the associated bursts of astrocytic Ca2+ activity might function to ensure an abundance of blood flow to the tissue in the aroused state, keeping the neurons metabolically prepared to handle additional auditory stimulations.

An important advantage of our volumetric imaging approach was in its ability to follow the spatial and temporal dynamics of dilations more comprehensively than in past studies. Thereby, we could define sites of origin and direction of the dilations as they progressed along the different vascular compartments. This approach was instrumental to our discovery of the central regulatory role of the pre-capillary sphincters, which was difficult to appreciate in previous work that imaged dilations at either the capillary or the PA level separately. Likewise, fast volumetric imaging was necessary to identify the presence of the local astrocyte Ca2+ activity that precedes the passage of dilations from the sphincters.

The spatial dynamics of dilations were investigated in previous work, but without reaching firm conclusions about their origin. In those studies, mostly performed in anesthetized mice, dilations had to be evoked by artificial stimulations. Some authors reported that they initiated at lower cortical levels ^51^ triggered by activity along the capillaries ^5,19,49,52^, whereas others that they initiated at upper levels, with the recruitment of the PA from the surface arteries ^53,54^.

This diversity might reflect the different methods used for evoking the dilatory responses in other studies ^21^ and the fact that a blood flow increase can be initiated by stimulation at any cortical lamina ^55^. In our study focusing on the PA-capillary junctions in awake mice, we observed a natural variety in the sites of origin of the dilations. Thanks to simultaneous fast imaging of all the relevant compartments, we could capture the specific directionality of these natural dilations, finding that some arrived from capillaries, other from PA, and others initiated at the junction level. Interestingly, a similar variety of originations was recently described in the barrel cortex of awake mice ^21,56^. This is not surprising because the two regions are similarly organized for blood flow regulation, with a large number of branching vessels ^57^, relays in the neuronal circuits, and projections from other cortical areas ^58^. In brain regions with simpler organization, the observed blood flow responses are more stereotyped ^19,59^.

The different positions in the auditory cortex at which the naturally occurring dilations originated must depend on differences in the modes by which blood flow regulation is recruited, which, in turn, may depend on the diversity of the neuronal circuit relays that project the sensory stimuli to the processing cortical region ^60-62^. In our analysis of tone-evoked dilations, we propose a classification based on the positioning of the dilations relative to the IOS response, which we considered to represent the core of the neuronal excitation. When the IOS response coincided with the area supplied by the PA, the region of neuronal excitation triggering the dilation most probably was along the PA inflow tract. In contrast, when the IOS response was in the vicinity but not exactly at the PA, the neuronal excitation likely occurred further down in the vascular tree and triggered the dilation around the capillary compartment (Figure 7a). We believe that this classification is roughly reliable, although it may not fully grasp the complexity of the phenomena giving rise to the dilation patterns that we observed. Additional factors may need to be considered. For example, the fact that the tonotopy has a more articulated nature than the IOS map, given its heterogeneity at the cellular level ^63,64^ with a diversity of neuronal activation patterns capable of triggering the NVC response. In addition, the fact that the sounds that induce the natural vascular responses are more complex than the three tones that we used here to evoke them, and complex sounds are known to often trigger tonotopic neuron responses in separate fields of the auditory cortex in parallel^65^.

**Figure 7.**
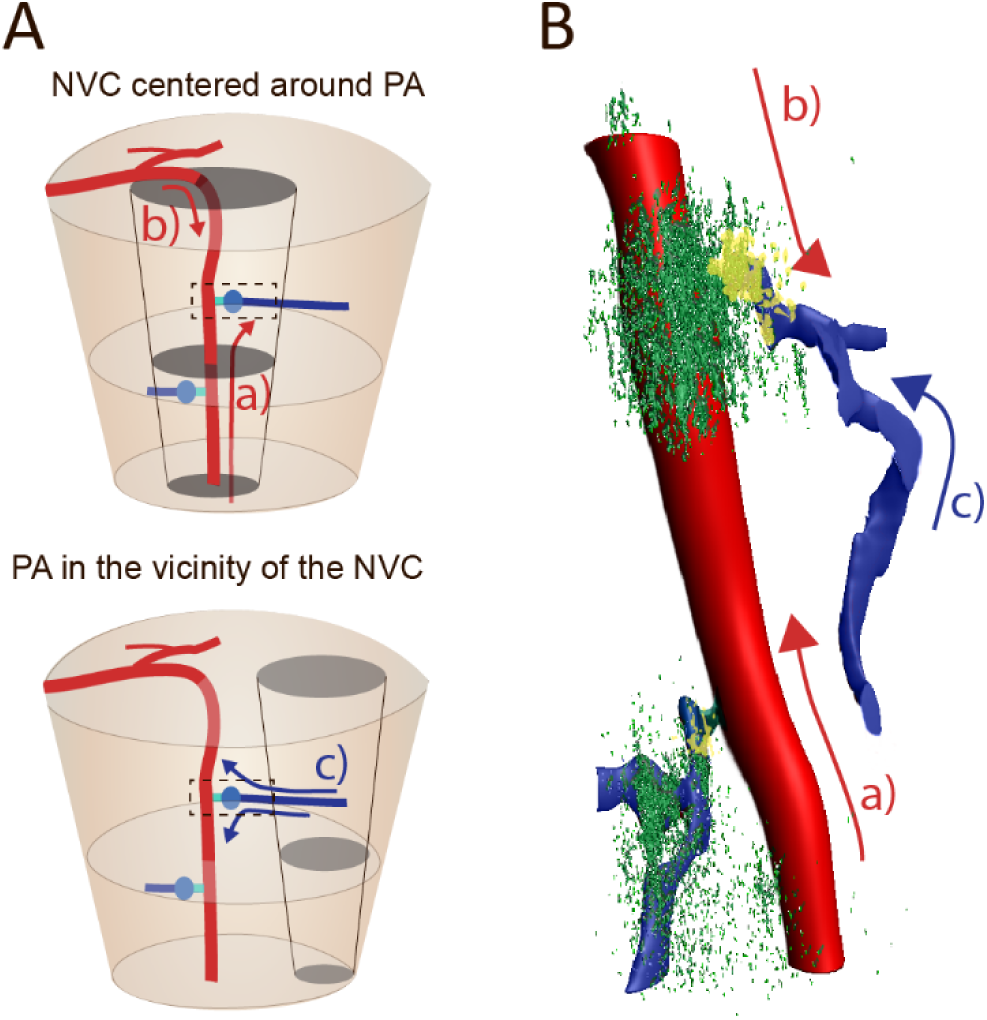
Schematic representation of the mode of progression of vascular dilations at upper cortical junctions under the control of astrocyte Ca2+ activity. A. Origin and direction of dilations at any given PA-capillary junction with respect to the areas of neuronal excitation. *Top*: case in which the excitation (*grey regions*) occurs exactly in cortical areas around the PA (*red*). In this situation, the NVC is either (a) initiated in the underlying layers by the excitation of layer IV-VI neurons or (b) arrives from the surface vessels due to widespread excitation. Either way, the dilation is perceived to arrive at the junction (*dashed-line rectangle*) from the PA (*red arrows*). *Bottom*: when the neuronal excitation (*grey regions*) occurs in regions close to, but not coincident with, the area directly supplied by the PA (*red*), the dilatory response arrives at the junction (*dashed-line rectangle*) from the capillary compartment (c, *blue arrow*) and will first recruit either the capillary (*blue*) or the pre-capillary sphincter (*cyan*). B. Illustration of how astrocyte Ca2+ activities at strategic locations regulate the spread of dilations across the sphincter between the PA and the 1^st^ order capillary. Astrocytes (*green*) regulate the progression of dilations with local and short-lasting Ca2+ activities in the end-feet enwrapping the pre-capillary sphincters (*yellow*). This regulation occurs independent if dilations arrive from the PA in deeper cortical layers (a, *red arrow*), or superficial layers (b, *red arrow),* or from activity along the capillaries (c, *blue arrow*), as detailed in panel A.

In addition to describing the varied origin of natural dilations in awake mice, our study describes, for the first time, the central role of the pre-capillary sphincters in determining by which extent these natural dilations spread between PA and capillary compartments under local control by astrocyte Ca2+ signaling. Pre-capillary sphincters have been well characterized anatomically only in recent years and only in the somatosensory cortex^6^. They have been described as a hemodynamic structural division between capillary and arterial blood flow, acting as a bottleneck opposing high resistance to blood flow ^9^. In view of their strategic interposition between PA and capillaries, sphincters in the upper cortical layers likely function to protect capillaries from high arterial pressures under baseline conditions. However, sphincters express α-smooth muscle actin in their mural cells, suggesting that they do not provide only passive flow resistance but can also actively contract. Indeed, previous experiments in anesthetized mice have shown that sphincters respond with significant changes in diameter to whisker stimulation^6^ or vasoactive agent infusion^8^ and thus could be involved in blood flow regulation during endogenous functional stimulation. The sphincters investigated here were not anatomically homogenous, for instance they differed in their level of indentation. However, contractile mural cells completely cover this part of the vascular inflow tract, thus all sphincters must be contractile ^6,37^ and their morphological differences may reflect mainly the extent of force in that contractility. Here, we provide the first direct evidence that pre-capillary sphincters play a physiological role during the auditory NVC response. Due to their strategic placement at the PA’s branching points, sphincters can control blood supply to larger parenchymal domains than downstream contractile capillary pericytes. Thus, a single PA connects to various capillaries that, in turn, branch out and supply blood flow to distinct sensory areas ^53,66^. Flow regulation at the PA’s branching points could optimize downstream blood delivery in register with neural activity ^67,68^ and be the base of the laminar activation pattern observed in human fMRI brain imaging ^69^. On the other hand, spread of dilations in the opposite direction, from capillaries to PA ^19,70^, can be equally important for pairing the increased local demands to global activation and contribute to the accurate and sufficient blood distribution essential for optimal brain function and health^5,36^. This upstream-directed regulation was reported to depend on endothelial K+ signaling^20^ triggered by TRPA1 or KIR channels stimulation directly on 1^st^ and higher-order capillaries ^71^, supported by another level of upstream-directed regulation at higher-order capillary branch points ^72^. While all the above observations align with our current results, they have provided only a fragmentary picture of the NVC-related dilation dynamics. Thanks to our 3D imaging approach and the large number of natural dilations that we recorded, we have been able to put pieces together and can propose the following interpretation of the observed dilation patterns: dilations arriving from the PA compartment may or may not involve the capillaries. The main factor determining their spread beyond the pre-capillary sphincter with increased blood delivery to the downstream capillary bed is most likely the position at which the neuronal activation driving the NVC response occurs (Figure 7b). If the activation localizes primarily to cortical layers deeper than the layer II/III junction, the dilation will generally involve just the PA. However, if the activation also concerns the upper cortical lamina, the downstream capillary region will receive additional local blood flow via expansion of the sphincter and 1^st^ order capillary. Likewise, a local neuronal activation can trigger dilations that initiate at the 1^st^ order capillary^5,19^, which will only involve the capillary branches. However, if the activation is part of a larger neuronal response, it will recruit additional blood from the surface vessels by spreading across the sphincter to increase delivery via the PA (Figure 7b).

Our study also describes for the first time the role exerted by astrocytes via local Ca2+ signaling in controlling the spread of dilations at the pre-capillary sphincters. The existence of the astrocyte control is supported by the observation that IP3R2KO mice, which lack part of the astrocyte Ca^2+^ signaling, display an altered NVC response, specifically a decreased capacity of dilations to engage the vascular compartments across the sphincters with reduced blood redistribution. ^325671^ Several past studies rejected a contribution of astrocyte signaling to NVC ^16-18^ because they failed to observe any changes in the blood flow responses in IP3R2KO mice. This may not be very surprising for the early studies in view of their experimental pitfalls, including having been performed in anesthetized animals. However, similar negative conclusions were drawn also in a recent study^17^ that was performed in conditions similar to ours’, i.e., in awake mice and considering multiple compartments, from PA to 4^th^ order capillaries. We do not contest the observations by Del Franco et al., as we could reproduce, using their analytical conditions, a lack of changes in some parameters of the dilation patterns in IP3R2KO mice (Sup. figure S9A). Instead, we highlight the important differences in the way experiments were conducted and data analyzed in their study compared to ours’ and which likely explain the discrepant conclusions. First, Del Franco et al., did not image all the vessel compartments at the junction simultaneously; second, they did not consider the pre-capillary sphincter region in their analysis, nor the site of origin of the dilations; third, they evoked dilations by stimulations that elicited widespread Ca2+ elevations in astrocytes not directly comparable to the local endogenous Ca2+ elevations that we observed just before the naturally occurring dilations, and, finally, they did not image the dilations along the z-axis. We could identify vascular abnormalities in IP3R2KO mice because we used a fast 3D imaging approach and focused on an aspect never investigated before, the dynamic progression of dilations across the sphincter. This led us to a twofold discovery: first, the recognition of the pre-capillary sphincter as a locus of physiological blood flow regulation, and second, the identification of a local astrocyte IP3R2-dependent Ca2+ activity controlling the sphincter’s contractility.

One of the problematic aspects in studying the functional roles of the astrocyte Ca^2+^ activity is its high level of complexity, involving a panoply of signals whose interpretation often remains enigmatic ^32,73^. Such signals go from highly visible, spatially large, and temporally long Ca2+ elevations, to more elusive, local, and fast Ca2+ transients. Considering that the Ca2+ signals relevant to the control of synaptic activity were identified as part of the local transients occurring at the level of peri-synaptic astrocytic processes ^32,74-76^, we expected that vascular-relevant signals would be mostly seen at astrocytic end-feet. Therefore, we focused attention on the Ca2+ events visible in the perivascular compartment, themselves quite heterogeneous ^15,77,78^. Aiming at identifying among them potential dilation-regulatory events, we recorded and analyzed the perivascular astrocyte Ca2+ dynamics having in mind how the extra blood flow was recruited in our experiments. Thus, we focused on the naturally-occurring vascular changes and 3D-imaged the peri-vascular regions around the pre-capillary sphincters that regulate blood flow to a specific cortical lamina. Since this approach had not been attempted before, we did not apply strict analytical criteria for defining the relevant astrocyte Ca2+ activity, rather defined a number of features that we expected such activity to have, like: (a) occurring in the end-feet within short distance from the vessel surface; (b) being time correlated to the dilations, i.e. shortly preceding them; and (c) appearing in association with all the dilations having a given pattern, i.e. originating at a given site and being able to cross the sphincter. As discussed, distinct dilation patterns reflect distinct types and loci of neuronal activation, suggesting that astrocytic Ca2+ activities related to distinct dilation patterns should also be diverse. Indeed, we observed pre-dilation end-foot astrocyte Ca2+ responses with different onset times and positions along the PA-capillary junction. Nonetheless, our “loose” analytical approach was strong enough to identify the NVC-relevant astrocyte Ca2+ activity, particularly because we sampled only physiological vascular events induced by auditory perception, either occurring naturally in the resting mice, or evoked by natural sounds. Our observation that the frequency of local astrocytic Ca2+ transients as well as the frequency of dilations progressing across the sphincter were both reduced in IP3R2KO mice, provide strong support to the relevance of the identified astrocytic Ca2+ activity to the NVC auditory response. Importantly, IP3R2KO is known to only partially reduce and not abolish astrocyte end-foot Ca2+ activity in the awake mouse ^45^, as we also observed for the pre-dilation Ca2+ events preceding natural (Figure 4) and tone-evoked dilations (Figure 6). Thus, it is possible that a more effective interference capable of abolishing any astrocyte end-foot Ca2+ activity might have revealed a stronger control by the astrocytes.

Some key aspects of the vascular regulation identified here remain to be addressed in future investigations. To start, the exact functional implications of the local control of blood distribution at pre-capillary junctions, by which dilations are enabled to expand their parenchymal territories of blood supply bidirectionally, as well as the extent of the functional consequences that alterations targeting this physiological mechanism can produce on brain function in pathological conditions. In this context, the fact that during several CNS diseases, astrocytes undergo morphological changes that alter or impede the contact between end-feet and blood vessels surface ^79^, implies noxious consequences for the regulatory functions here identified, that could likely contribute to the disturbed blood flow seen in such pathologies ^1^, notably in those accompanied by cognitive decline ^80^. Another aspect to be clarified in the future is the additional impact of the large peri-vascular astrocytic Ca2+ activities that we saw invading several end-feet during movements or arousal states of the mice. Considering the astrocyte Ca2+ control at the pre-capillary junction in the resting mouse, it is intriguing that when we saw large Ca2+ events during mouse movements, we also observed an increase in multicompartment dilations (Figure 2F). It remains to be defined how exactly these astrocyte phenomena influence NVC during motion and/or changes in brain states involving activation of secondary pathways and/or neuromodulator effects that overlap with the primary auditory excitation. ^81,82^. Understanding these relations will lead to a more comprehensive understanding of how astrocytes contribute to keeping healthy and adequate levels of substrate and oxygen in the brain of behaving mice, and how these levels could be altered under pathological brain conditions. In this context, deciphering the mechanism(s) triggering the relevant Ca2+ activity in astrocyte end-feet and the subsequent changes in sphincter responsivity would be important. Are the local astrocyte Ca2+ elevations secondary to neuronal activation and synaptic transmission ^32,74^? How are they related to larger Ca2+ elevations triggered by specific brain states and changes in noradrenaline levels ^39,43^? Several mechanisms have been described by which astrocytes can sense synaptic activity locally and generate a Ca2+ increase response ^83^. To start, via activation of Gq-GPCRs that signal via IP3 like mGluR5 or P2Y1R ^26,75,84^ and whose effects could be attenuated in IP3R2KO. However, also ionotropic receptors like P2X1 and P2X5 can produce direct or indirect intracellular Ca2+ influx in astrocytes^11,14^. Other mechanisms capable of producing astrocyte Ca2+ elevations independent of IP3R2 receptors could involve plasma membrane Ca2+ influx channels like TRPA1 ^85^ or TRPV4 ^86^, or the activity of Na+/Ca2+ exchangers following neurotransmitter uptake ^87^. Activation of TRPA1 channels in endothelial cells^71^ could also initiate a local Ca2+ response in astrocytes by triggering K+ release onto their end-feet.

While our study leaves these important questions open, we believe that its identification of the peculiar control exerted by astrocytes at the sphincter, a critical point of blood flow regulation in the vascular tree, significantly advances understanding of the astrocytic contribution to NVC.

## Materials and Methods

### Animals

Most experiments in this study utilized mice induced to genetically express the Ca2+ sensitive indicator GCaMP6f selectively in the cytosol of astrocytes based on the GFAP promotor. This was obtained by crossing two transgenic mouse lines, a lox-STOP-lox-cytosolic-GCaMP6f (purchased from The Jackson Laboratory http://www.jax.org, JAX 024105 (*B6;-Gt(ROSA)26Sortm95.1(CAG-GCaMP6f)Hze/J*), and *hGFAPCreERT2* mice ^88^ obtained from Prof. F. Kirchhoff, University of Saarland, Germany, as previously described ^32^. A subgroup of these *GFAPCreERT2xGCAMP6f* mice was then crossed with the *Itpr2KO* mouse strain ^89^ that carries a constitutive deletion of IP3R2 expressed in the ER and displays conspicuous reduction in astrocytic Ca^2+^ activity ^16^. We obtained this strain from the laboratory of prof. Ju Chen of University of California San Diego, La Jolla, California, US. We used only IP3R2 ko/ko homozygotes in this study. Though IP3R2 contributes importantly to astrocytic Ca^2+^ activity, especially in Ca^2+^ mediated Ca2+ release from the ER in soma and large processes, not all the astrocyte Ca^2+^ activities depend on this receptor, and only the largest are abolished in the knockout ^90^. The mice were treated with Tamoxifen at 8 weeks of age to induce *Cre* recombination and trigger GCaMP6f expression in astrocytes. Tamoxifen (Sigma-Aldrich) was dissolved in corn oil (10mg/ml) and administered i.p. (0.1 ml/10 g body weight/day) for 5 days prior to the chronic cranial window surgery. The mice had surgery when they were 8 weeks old and, after a 4-week period of recovery and training, were included in the experiments between 12-28 weeks of age. Two 12-16 weeks-old *GFAP-EGFP* mice ^38^ were used for morphological investigations. All experiments and procedures were conducted under license and according to regulations of the Cantonal Veterinary Offices of Vaud (Switzerland).

### Chronic cranial window preparation

Chronic cranial windows were established as previously described ^32^. To target the auditory cortex, the window was centered around -3mm anterior-posterior (AP) and 5 mm mediolateral (ML) from bregma, right hemisphere. The surgical procedure was done under isoflurane 1.5% anesthesia supplemented with a Carprofen injection (5 mg/kg, s.c.) and local anesthesia in the form of Lidocaine (0.2 %, s.c.) under the scalp. Animals were kept warm on a temperature-controlled heat blanket during the procedure, and their eyes were protected against dehydration with viscotears. Hair was removed, and Betadine used to sterilize the skin prior to the first incision. A square hole 3x3 mm was drilled in the bone and, leaving the dura intact, was covered with a fitting glass coverslip (thickness #1), which was then glued to the cranium. A L-shaped metal plate used to head-restrain the mice during the imaging was fixed to the skull by glue and dental cement. Analgesics were given for three days after surgery (Paracetamol 125 mg dissolved in 250 ml water).

### Electromyography (EMG)

We recorded animal movements by electromyography (EMG)^91,92^. To achieve this, two electrodes were implanted within the neck muscles of the mouse after the chronic cranial window preparation and connected to two micro sockets. During *in vivo* imaging, the sockets were connected to a preamplifier and a Molecular Devices digitizer (Digidata 1440A). The signal was recorded using pClamp and then filtered (50 Hz filter) using a custom-made Matlab script. The onset of each recording was triggered by a digital signal from the Bruker microscope upon initiation of each two-photon imaging sequence (see “*3D awake imaging”* section). The recorded EMG signal was first filtered using a 50 Hz notch filter. Then the filtered recording was normalized to standard deviation values, using the mean and SD values from the entire 90-120 seconds acquisition period. Timepoints in which the normalized signal reached >5SD were identified, grouped together and considered part of the same movement event if they were <500 ms apart.

### Fluorophores

To visualize the lumen of the blood vessels, just prior to the imaging session we introduced in the circulation by tail-vein injection a dextran conjugate with Texas red 70.000 MW (2 % in 0.9 % NaCl sterile solution, 100 µl bolus I.V.). The vascular dye was of a size that impedes extrusion to the tissue surrounding the vessel. Hence, it was ideally suited to highlight the lumen of the blood vessels and signal diameter changes ^14,93^.

### 3D Awake Imaging

Before imaging, the mouse was habituated to human handling for two days and gradually introduced to being head-restraint under the microscope and then thoroughly trained for an additional five days. The head of the animal was restrained by a custom-made system in which a metal plate attached to the head mount of the mouse was fastened with a screw to a matching metal bar. During imaging, the mouse sat on an air-supported, freely floating ball that acted as a spherical treadmill, allowing the animal to run when it wanted. The head of the mouse was slightly tilted, and the objective was at a 40° angle from the vertical position to allow perpendicular imaging through the glass covering the auditory cortex. Two-photon imaging was done with a Bruker *in vivo* Investigator system (Bruker Nano Surfaces Division, Madison, WI, USA) equipped with an 8kHz resonant galvanometer scanner, coupled to a MaiTai eHP DS laser (Spectra-physics, Milpitas, CA, USA) with 70 fs pulse duration, tuned to 920 nm. Negative dispersion was optimized for each wavelength, and laser power was rapidly modulated by Pockels cells. A 20x LUMPFL60X W/IR-2 NA 0.9 Olympus objective was used. Emission from red and green channels was separated by a dichroic beam splitter (565 nm Long pass) allowing shorter wavelengths to reach a 520/540m band pass filter before a GaAsP detector (520 nm, optimum for green emission) and longer wavelengths to reach a 610/675 nm band pass filter before another GaAsP detector (610 nm, optimum for red emission). The combination of a resonant scanner with a piezoelectric actuator made high-speed 3D imaging possible, while the highly sensitive GaAsP detectors with negative dispersion allowed applying minimal laser dose to the tissue. The laser power varied during experiments depending on the depth of the focus, but it was kept below 7 mW and measured continuously with a power meter. These imaging settings were previously thoroughly tested to ensure their non-toxicity to the tissue, even at high-speed image acquisitions ^32^. The anatomical localization of PA and 1^st^ order branch points defined the depths at which the imaging was performed. All the junctional sites except one were located ≥80 µm below the brain surface, with a maximal depth of 328 µm. Two different volumetric imaging approaches were applied: in one, imaging was performed in small volumes with stacks centered around the PA-capillary junction, and in the other, in larger volumes with stacks following the PA up and down the upper cortical layers. More specifically, in the first approach, the 3D-FOV covered an imaging volume of 56-75 µm x 15-44 µm x 21-35 µm (x,y,z), the acquisition speed was 200 Hz and the optical-zoom 8x, with pixel size of 0.29-0.59 µm lateral resolution and 1 µm axial resolution. In the second approach, the 3D-FOV covered an imaging volume of 75-121 µm x 75-112 µm x 190-280 µm (x,y,z), the acquisition speed was 200 Hz and the optical-zoom 8x, with pixel size of 0.29-0.94 µm lateral resolution and 10 µm (and in one case 20 µm) axial resolution. The scanning rate per 3D stack was 8-10 Hz. A maximum of 20 imaging sequences, 90-120 seconds-long (800-1000 stacks in total), were taken with 1-5 min breaks between them, depending on the mouse behavior.

### Auditory stimulations

In this set of experiments, mice were subjected to a single sound stimulation during each of the imaging sequences. Sounds were produced by an Electrostatic loudspeaker (Tucker-Davis Technologies, Inc.) placed 30cm from the left ear of the mouse, connected to the pClamp digitizer. The latter was programmed to produce a single 10 s-long, 1 Hz auditory stimulation consisting of 10 repetitions of an individual tone, each one lasting 500ms, that was started 30 s after the initiation of the imaging sequence. Three different tone frequencies were used as stimuli: 3 kHz, 20 kHz, or 30 kHz. Each tone stimulation was repeated during three different imaging sessions.

### Intrinsic optical signaling (IOS)

In the auditory stimulation experiments, before performing two-photon imaging, we used intrinsic optical signaling for obtaining a tonicity map of the auditory cortex. Using the 4× objective of our microscope, we could include the entire cranial window in the field of view. The light source was filtered with a green light excitation filter (532 nm). This wavelength is equally scattered by oxygenated and deoxygenated blood, so the reduction in the emitted light reflects changes in total blood volume ^94^. We sampled images at 10 Hz and compared 10 s periods before and during tone stimulation, repeating comparisons for all the tone frequencies. We calculated the difference light scattering before and during stimulations ^95^ to detect the position of the largest hemodynamic response to each of the different tone stimulations.

### Mouse pupil imaging

In a sub-group of the two-photon imaging experiments, we performed pupil imaging using a high-resolution, fast-speed IR camera centered on the eye of the mouse. The camera was a Dalsa Genie Nano, run with Sapera LT and CamExpert software. Pupil imaging was started by a digital signal from the Bruker microscope upon initiation of each imaging sequence. A bandpass filter (850 nm) was placed before the objective to exclude visible light, only permitting visualization of the mouse pupil by the reflected IR light. This enabled us to follow the pupil contractions and dilations during two-photon imaging and to quantify them using a simple custom-made MATLAB script. Periods involving pupil dilations (expansions) were detected based on the data z-score and defined as pupil enlargements >2SD with respect to the average size of the pupil during the whole imaging period. The timings of pupil expansions were compared to those of astrocytic Ca2+ elevations averaged across the entire FOV, roughly measuring large astrocytic Ca2+ events. Periods in which both measurements reached values >2SD were defined as periods of occurrence of the two phenomena in overlap (Suppl. Fig S2D).

### 3D imaging data analysis : general aspects

The data obtained via 3D two-photon imaging were analyzed with custom-made MATLAB (Matworks, 2019b version) scripts and visualized using both ImageJ and Imaris 8.2 software (Bitplane). The analysis consisted of several steps. First, with the script “VesSegCreateMasks.m”, each z-level (or focal plane) in the 3D stack was considered as an independent 2D image (Sup. Figure S1A and B). In each of these images, the user manually defined the position of the different vessel compartments and drew a rectangular region around each of them (Sup. Figure S1C). In the “VesSegRoiData.m” script, these selections were then applied to the whole 3D timeseries, i.e., the rectangular regions were automatically used in each focal plane, so that each region contained and roughly outlined a specific vessel compartment (Sup. Figure S1D and S3). Then, the vessel compartments and the related astrocyte end-feet structures were automatically detected within each rectangle, their position found throughout the imaging sequence, and the areas calculated at each of the z-levels. Detection of the vessel structures was done first by normalizing all the pixel intensities in the red channel within the selected rectangular region during the entire imaging sequence and by calculating the z-score of each pixel. Then, the area corresponding to the structure within the rectangular region was masked at each time point using a gaussian filter (imgaussfilt.mat, with the variable sigma decreasing from 8 to 5 with the increase of the imaging depth). The mask’s area was defined by the sum of all the red channel pixels in the rectangle that had significant fluorescence level (>2SD) for all the time points in the timeseries (Sup. Figure S1E). Movements were corrected based on the position of the centromere of the specific mask in the rectangular region under analysis compared to the centromere of the average mask from the entire imaging sequence, and the operation repeated to reposition the rectangular region for each time point. To detect whether a vessel structure was contained within the rectangular region, we applied a size threshold and established that the vessel structure within the rectangular region should cover >8% of the region’s area to be retained for further analysis (bwareaopen.mat). In cases in which movements of the mouse caused exit of the vessel structure from the 3D-FOV at a given time point and in which repositioning of the rectangular region could not recover the structure, the timepoint was deleted from the entire 3D+t stack and the data thus excluded from the study. Such a deletion was sometimes necessary during substantial movements or locomotion of the mice. The GCaMP6f signal in the green channel within the rectangular selection was normalized in the same way as done for the red channel. From this data, end-feet structures were defined for each imaging plane as structures departing from the edges of the vessel structure and occupying the first 2 µm external to such edges, which gave a 3D annulus ROI around the vascular structure (Sup. Figure S1F). These ROIs would then comprise all the Ca2+ activity occurring in astrocytic structures within 2 µm from the vessel structure, regardless of the astrocyte to which the end-foot structure belonged. The ROI’s width of 2 µm was chosen based on previous EM descriptions of the continuity and thickness of the end-foot layer around CNS vessels ^96^. In fact, the end-feet thickness in EM is ≤1 µm, but we added a conservative +1 µm margin to ensure its complete inclusion taking into account the scattering of the emitted light during two-photon imaging. This ROI was divided into four quadrants that enabled repositioning the ROI in relation to the vessel position and its constriction/dilation activity along the x and y axes (Sup. Figure S1G-I). Thus, first, the position of the rectangular region was adjusted according to the detected movements, and then the end-feet ROIs were moved in relation to the position of the vessel mask.

### Detection of dilation events and patterns

The following analytical approach was utilized both for the imaging experiments in small volumes at PA-capillary junctions and for those in larger volumes along PAs and both for natural and tone-evoked dilations. At first, the vascular structures within the 3D-FOV were classified as PA, sphincter, bulb, or capillary compartments utilizing the 3D stacks reconstructed from the ensemble of the rectangular regions manually drawn for each 2D image at a given z-level (Sup. Figure S1C and S3B). To note, some of the regions in the stacks were empty, i.e. lacked their structure at specific z-levels, especially in stacks covering large volumes with long steps between z-levels. Next, the spatial-temporal pattern of each dilation event was defined using the custom written “VesSegActivitySearch.m” script. For each vessel compartment, the vessel area at each z-level was normalized to SD values (Sup. Figure S3C), which were calculated taking periods without movements as baseline. By combining the relative changes in area at any z and t data point, we created a 3D image of the vascular dynamics in time and depth (Sup. Figure S3D). This z-t rendering of the dilation in 3D was then normalized (A-mean(A) /std(A)) considering all the time points in the imaging sequence (where A = area). Finally, to define dilation periods that extended three-dimensionally along the vessel structures, a 2SD threshold was used to detect dilations that occurred in several z-levels, prior to a gaussian blurring (imgaussfilt.mat, sigma=2) followed by removal of ambiguous dilation events that were either highly local or very short (bwareaopen.mat, filter: ¼ x z-levels x imaging frequency) (Sup. Figure S3D). Via this procedure, the vascular activity (Sup. Figure S3E) was simplified and binarized into periods of either dilation (1) or no dilation (0). Binarization was performed for each vessel compartment during the whole acquisition period (Sup. Figure S3F). By comparing the information from the different compartments, we could evaluate if the onset of each dilation event occurred within or outside the 3D-FOV. Moreover, we could establish which vessel compartment dilated first and when, as well as the number of other vessel compartments that participated in the dilation and the onset time of the dilation in each compartment relative to the initial one (Sup. Figure S3G). Based on this information, dilations involving more than one compartment were considered a single event when they occurred in temporal continuity in the different compartments, whereas they were considered separate events when they were spaced by >500 ms intervals. In tone stimulation experiments, dilations occurring within the tone stimulation period were considered to be evoked. The delay from stimulation to dilations’ detection in the FOV ranged between 1.9 and 3.3 s depending on the dilations’ direction, in line with the commonly reported delay of evoked NVC responses^37^. We excluded movement-related events as components of such dilations but cannot fully exclude contribution by other factors such as coincident natural sounds.

### 3D imaging at the PA-capillary junction

#### - Detection and analysis of pre-dilation astrocytic 3D Ca2+ activity

Astrocytic pre-dilation Ca2+ activity was investigated in small volume 3D imaging stacks centered around the sphincter based on the GCaMP6f green fluorescence signals. Once we had quantified the timing and pattern of vessel dilations based on the Texas red fluorescence in the vessel lumen (see “*Detection of dilation events and patterns”* section and Sup. Figure S5A-B), the onset of each dilation event displaying a given pattern was used to identify Ca2+ activity in astrocytic end-feet that had the specific feature of appearing recurrently with a defined temporal relation with the dilation event (Sup. Figure S5C). For this, we used the custom written VesSegPreDilCa.m script. To start, we considered the average astrocytic Ca2+ activity present in each end-foot ROI around a vessel compartment at each z-level of the 3D-FOV stack and normalized it to SD values, defining the baseline mean value and the SD values from the periods without movements (F-meanF_baseline_)/stdF_baseline_). As done for the vessel dilation data, this normalized single-plane information was combined in a z-t matrix, obtaining a 3D+t reconstruction of the average Ca2+ changes in the astrocytic end-feet ROIs shown as changes along the z-axis over time at each x-y imaging level. We then temporally aligned this map of the 3D+t astrocyte Ca2+ changes to the onset time of each dilation, considering specifically a 3-seconds period including the 2 seconds preceding the dilation event and the first second after its start, the latter to avoid abrupt cut of Ca2+ dynamics started in the pre-dilation period but still ongoing at dilation onset (Sup. Figure S5D). The 2-seconds pre-dilation period was chosen based on the expected delay of vascular responses from neuronal activation according to past NVC studies ^15,19,49^ and our own previous observations ^23^. Noteworthy, some of the naturally occurring dilations here investigated started outside the 3D-FOV, so for calculating pre-dilation delays we could rely just on their observable timings, i.e. when they entered our 3D-FOV. The astrocytic Ca2+ activity was then normalized along all the z-levels over the entire 3 s selected period. Within this time frame, all the z-axis locations and timings in which Ca2+ events occurred were identified by using a >2SD threshold for defining a Ca2+ increase, gaussian filtering with a 0.5 sigma gaussian blur (imgaussfilt.mat), and excluding events smaller than 3 µm x 100ms (bwareaopen.mat). The result was a binary mask in z-t defining the temporal and spatial position of each of the astrocytic end-foot Ca2+ activities in relation to each dilation event. We then regrouped all the astrocyte Ca2+ masks that were associated to dilations with the same pattern, i.e. same onset location and same vessel compartments involved. The regrouping of the masks from all the imaging sessions of a given experiment allowed their comparative analysis. We found that the pooled Ca2+ masks contained some overlapping Ca2+ activity, i.e. activity that occurred in all of them at the same time and 3D location. We defined the recurrent regions as the VOIs (voxel of interests) putatively involved in the pre-dilation astrocytic Ca2+ activity. We then came back to each specific map of Ca2+ activity associated with each individual dilation event, overlaid the template VOIs map to it, and checked if the individual map showed dilation-related astrocyte end-foot Ca2+ activity at the z-level and time identified by the VOI (Sup. Figure S5E). The size, timing and duration of these average Ca2+ signals were then used to describe the spatial and temporal extent of the pre-dilation Ca2+ signals (Figure 3C). We considered that a dilation was preceded by this type of pre-dilation Ca2+ activity when the Ca2+ activity overlapped with the average z-t VOI at that specific position in 3D-t. Finally, we investigated how recurrently each putative pre-dilation Ca2+ activity was detected prior to dilations regrouped by the same pattern (Sup. Figure S5F). As inclusion criterion we considered the frequency of occurrence, with a margin defined by the number of dilations belonging to a given pool and using a 0.1 confidence limit (1-(((N/2)-((sqrt(N)/2)x1.282)/N)). Hence, for example, if a pre-dilation Ca2+ signal occurred in 59 % of N=50 dilations of the same type, we considered it to be a recurring event related to that dilation pattern because it was within the margin of 41 %. In the case we had only N=10 dilations, the event would have been considered recurring if it was seen in 70 % of the dilations due to an error margin of 30 % (Sup. Figure S5F). This criterion was adopted to avoid positive bias towards dilation patterns that occurred rarely. This means, however, that our criterion was more restrictive in the experiments on tone-evoked dilations because in those experiments we observed a much lower number of dilation events than in experiments on naturally-occurring dilations. Nonetheless, the number of pre-dilation Ca2+ events that we identified in the tone-evoked dilation experiments was in line with the number in experiments on naturally-occurring dilations (Sup. Figure S10D and S10E).

#### - Detection of large astrocytic 3D Ca2+ activity

In small volume 3D-FOVs at the PA-capillary junction we also identified periods with large pre-dilation astrocytic Ca2+ activity during resting periods. Analysis was performed on the z-t projection of the end-foot Ca2+ activity ROIs as for the small Ca2+ activity. This was done because our pre-dilation Ca2+ detection approach, described in the previous section, would not distinguish small from large activities. To tease apart the two types of events, we used the z-t matrix plot and defined “large astrocytic Ca2+ activity” as periods of extensive activity where Ca2+ increased >2SD above the baseline value in non-movement periods and spread through the z-layers to include the entire depth of the z-stack in >75 % of the end-feet ROIs. The dilations that occurred within these periods were then excluded in the following analysis of pre-dilation Ca2+ events.

### 3D imaging along the PA

#### - Detection and analysis of astrocytic Ca2+ activity

In experiments in which we scanned large volumes along the PA, the astrocytic end-feet structures were imaged in focal planes 10-20 µm apart along the z-axis (Figure 1C). Thus, the astrocyte structures present in the 2 µm annulus created along the vessel for each focal plane were treated as individual entities, and their activities averaged within the corresponding focal plane and analyzed separately. In this type of analysis, astrocytic Ca2+ activities were defined as peaks standing out of the mean baseline fluorescent signal from all the end-feet surrounding the PA at the given z-level. Peaks were detected starting from the normalized ((x-mean)/sd), smoothed (`sgolayfilt.mat`, order value=10) mean baseline signal using the Matlab command `findpeaks.mat.`The following parameters were used: `MinPeakHeight` = 1 SD of the smoothed data; `’MinPeakProminence’`= 0.8 SD of the smoothed data, `MinPeakWidth` =1 second, and w`idthReference=halfheight.` After detection of the start and end times of the peaks, which correspond to the times in which the smoothed fluorescence data went, respectively, above and below the mean baseline value, we used the non-smoothed normalized data to verify that the signal prominence was >2 SD. Finally, we compared the spatial-temporal characteristics of the Ca2+ events determined in each focal plane with those in other focal planes and when we found that they overlapped, we classified the events in different planes as a single 3D Ca2+ activity, quantifying its z-spread (µm) and duration (s). The time periods showing Ca2+ events in end-feet along PAs were analyzed with regards to whether they overlapped with periods of animal movement or with PA dilation events to evaluate if correlations existed between the phenomena.

### Statistics

Data are presented as mean ± 95% confidence limit (C.L., 1.96xSEM), unless otherwise stated. For all statistical analyses, OriginPro 2018b (OriginLab, Northampton, MA), Matlab 2019b, and Excel (Microsoft Office 2016) software was used. Statistical analyses were performed per 3D-FOV. For experiments on naturally-occurring dilations involving imaging small volumes at PA-capillary junctions, we analyzed 20 FOVs in 6 WT mice and 14 FOVs in 4 IP3R2KO mice, respectively. In each individual statistical test, we provided information on the number of dilations analyzed/3D-FOV in terms of mean dilations/experiment ± 95 % C.L.. For experiments involving imaging large volumes along the PA, we analyzed 8 FOVs in 6 WT mice and, in view of the reduced number of FOVs, we gave just the N value corresponding to the total number of dilations observed. In terms of statistical analyses, initially we compared the distribution of two data groups using graphical quantile-quantile plots (Q-Q plots), i.e. comparing the quantiles of the two groups: when the distribution was normal, a parametric two-tailed t-test was used considering whether data were paired or not. When the two data groups could not be considered independent or did not have normal distribution, the data were analyzed using non-parametric tests: the Wilcoxon signed-rank test was used for paired data, and the Mann-Whitney test for non-paired data. In data sets with several groups, before comparing individual points from two groups, to exclude group effects, we initially performed the Kruskal-Wallis ANOVA (KWA) for non-parametric data, and the two-way ANOVA for normally distributed data. If the data points in a group could not be considered to be independent, as in the case of vessel compartment dilations, we initially used the Friedmans ANOVA test instead. KWA and Friedmans ANOVA tests provide both a p-value and a X^2^ value. The latter was used for defining the level of difference between the compared groups. In analyses that required a multiple comparisons test, such as ANOVA or KWA, the values from the final analysis were adjusted with the Holm correction (similar to Holm-Bonferroni) on both parametric and non-parametric data. This correction depends on the number of comparisons in the data group and the adjustment is performed in a ranked way depending on the p values: the smallest p-value gets the strongest adjustment (multiplied by the number of comparisons), while the following increasing p-values are adjusted depending on the number of p-values already adjusted according to: P_original_x(N_comparisons_-(counts of P< P_original_))=P_corrected_. Data were considered significantly different when the p-value, corrected or non-corrected, was *<0.05, but additional levels of significance were marked with **<0.01 and *** <0.005.

### Data availability

All scripts forming the code used in this study together with data examples to verify code function are provided with the submitted manuscript in a zipped folder together with a read-me instruction on how to run the analysis. All other data can be made available from the author (BLL) upon reasonable request and will be uploaded to a public repository upon publication of the study. A space in the public repository Zenodo has been reserved for this data under the doi: 10.5281/zenodo.15224578.

## Acknowledgments

We would like to thank Giovanni Carriero and Erika Bindocci for their advice and support during the project; Tania Barkat, for her advice on setting up the auditory stimulations; and Martin Lauritzen and Søren Grubb for valuable scientific discussions. This work was supported by ERC advanced “Astromnesis”, SNSF 31003A-173124, and SNSF 31003B-201276 grants to AV and by postdoc fellowships R210-2015-3320 and R265-2017-4437 to BLL from The Lundbeck Foundation, Denmark.

## Author contributions

BLL designed and performed all experiments, custom wrote all matlab analysis tools, performed the analysis and statistics, designed figures and wrote the manuscript. A.V. supervised the project, defined strategy, discussed experimental design and analysis, designed figures and wrote the manuscript.

## Supplemental Figures

**Sup. Figure S1.**
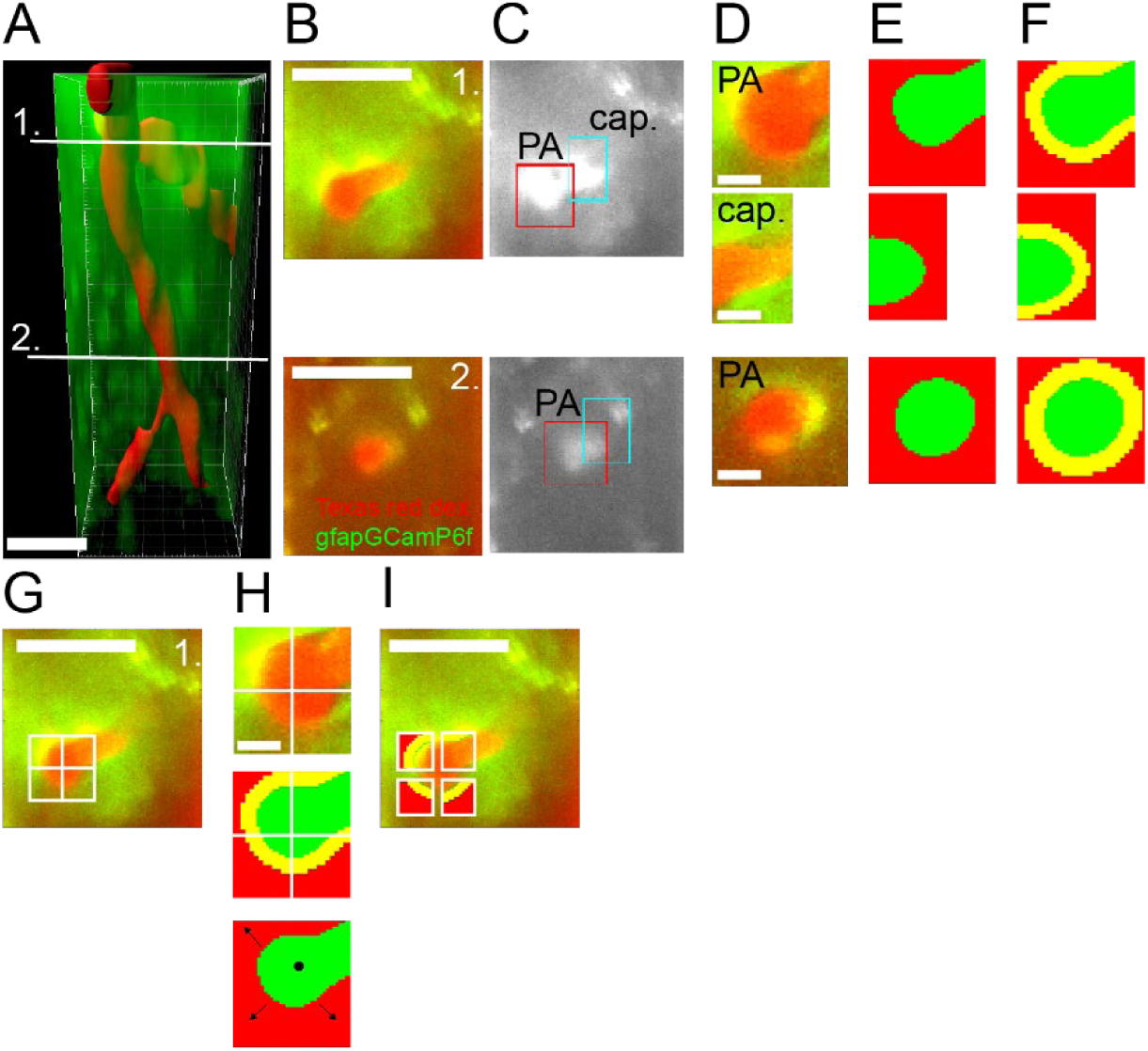
Method to detect vessel position and astrocyte end-feet ROIs within manually defined vessel segments in a 3D-FOV. A) 3D reconstruction of a PA with two 1^st^ order capillary branches taken 40-240 µm below the cortical surface. The image represents the cumulative average of a 3D-FOV (x= 75.5 µm, y= 75.5 µm, z=200 µm) timeseries (104 s) taken from the auditory cortex of a Texas red-dextran injected *GFAPCreERT2:GCaMP6f* mouse. *Red*: Masked vessel lumen. *Green*: cumulative GCaMP6f signal in astrocyte structures. *White lines* indicate the position of the two z-levels shown in B-F. Scalebar: 30 µm. B) Two 2D x-y images from the cumulative averaged x-y-z stack in A representing individual scanned focal planes at z-levels 1 and 2 respectively. *Red*: Texas red-dextran in the vessel lumen. *Green*: GCaMP6f signal in astrocyte structures. Scalebar: 30 µm. C) Rectangular regions framing relevant vessel segments were manually defined at the two z-levels shown in A-B. In each experiment, such rectangular regions were defined at each z-level for each 3D-t scan performed. Their manual definition was done consistently from one z-level to the other to follow the same vessel between layers. *Red rectangle*: PA segment; *blue rectangle*: capillary segment. D) Average fluorescence signals in the two cropped rectangular regions defined in C. The regions contain both the Texas red-dextran signal, outlining the specific vessel segments shown in C, and the GCaMP6f signal coming from the astrocyte structures surrounding the vessel segments. Scalebar: 5 µm. E) The area of each vessel segment (here depicted in *green*) in the cropped rectangles was automatically determined by masking the normalized, gaussian filtered images using a custom-written software (see Methods). F) End-feet structures (here depicted in *yellow*) were automatically defined for each cropped rectangle as 2 um-thick annular ROIs around the vessel structures in continuity with the vessel’s edges. G) Same rectangular crop from z-level 1 shown in C-F presenting the average red and green fluorescence from a Texas red-dextran injected *GFAPCreERT2:GCaMP6f* mouse. The cropped image captures the PA before dilation and was divided into four quarters to allow repositioning relative to changes in vessel diameter. Scalebar: 30 µm. H) Following dilation, the quarters in the rectangular crop shown in G were moved relative to the positions of the PA centromere (*black point*) and of the borders of the vessel lumen (*arrows*) to adapt to the increased vessel volume, as described in Methods “*3D imaging data analysis*”. Scalebar: 5 µm. I) The end-feet ROIs in the quarters of the cropped rectangle are repositioned following dilation relative to the new vessel position and diameter change. Only after this operation, measurements of astrocytic end-feet Ca2+ activity were made. Scalebar: 30 µm.

**Sup. Figure S2.**
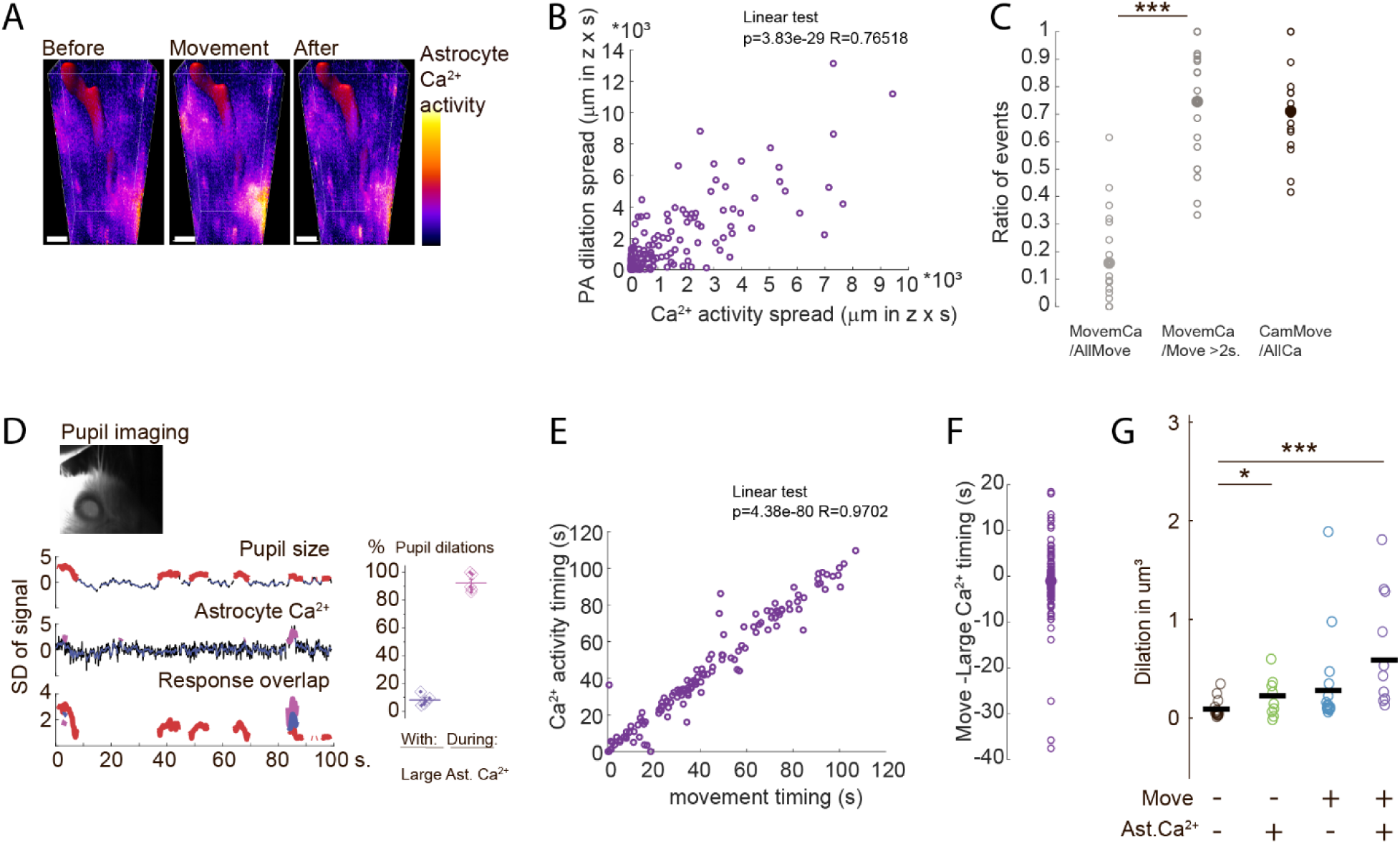
Correlation between movements, large astrocytic Ca2+ events, and dilations. A. Astrocyte Ca2+ levels measured as cumulative average of the GFAP-GCaMP6f signal (*pseudocolor*) in a large-volume 3D-FOV along the PA during 5 s periods when the animal was at rest before moving (l*eft*), when it moved (*middle*) and when it was at rest again after movement (*right*). *Red*: Masked vessel lumen. Cortical depth 40-240 µm. Scalebar: 20 µm. B. Linear correlation between the spatial spread (µm in z x s) of astrocyte Ca2+ activities and of PA dilations during movements (N= 145 dilations events, 8 PAs, in 6 mice), measured using 10-20 µm steps in z-stacks covering 100-280 µm of the depth of the cortex beginning from 50-100 µm below the surface (Pearson linear correlation p=3.9x10^-29^, R=0.76). C. Proportion of cases in which large astrocyte Ca2+ activity and movement of the mouse overlap during each experiment, relative to the total number of: movements (*left*), long movements (>2s, *middle*), or large Ca2+ activities (*right*). Considering all movements, they were accompanied by astrocytic Ca2+ activity in only some cases (*left*); however, considering just long-duration movements, the astrocytic Ca2+ activity was present in more cases (*middle*: Mann-Whitney test, p=0.0007, 11.1±4.6 Ca2+ events and 54.0±24.5 movement events in N=20 FOV in 6 mice). Considering the cases of large astrocytic Ca2+ activities, they appeared in association with a movement (of any duration) in 71.1±7.5% of cases. D. *Top*: Mouse pupil diameter changes recorded during two-photon imaging sessions. *Bottom, left*: example of simultaneous traces of pupil size (*top*), average astrocytic Ca2+ activities (*middle*) and their overlay (*bottom*). Pupil dilations periods (>2SD) are shown in *red*. Astrocyte Ca2+ activities periods (>2SD) are shown in *purple*. The overlay shows in *blue* the overlapping periods. *Bottom, right*: percentage of pupil dilations accompanied by a large astrocyte Ca2+ activity (*left*) and, vice versa, percentage of large astrocytic Ca2+ activities occurring in overlap with a pupil dilation (*right*). (N=95 astrocyte Ca2+ activities and N=450 pupil dilations in 5 mice). E. Linear correlation between the onset time of the astrocyte Ca2+ activity and the onset time of the animal movements considering all those movements associated with a large Ca2+ activity. N = 129 events from 20 FOVs. (Pearson linear correlation, p=4.37x10^-8^ R=0.97). F. Onset time of movements relative to onset time of Ca2+ activity in the same event population as in E (129 events from 20 FOVs). Movements initiated on average 1.03±1.25s before the astrocytic Ca2+ activity. G. Average vessel lumen expansion (µm^3^) during dilation events per FOV in different conditions: at rest (*black*: 162±31.2 dilations/FOV); at rest in the presence of a large astrocyte Ca2+ activity (*green*: 4.2±2.0 dilations/FOV); during animal movement without large astrocyte Ca2+ activity (*azure*: 47.6±22.1 dilations/FOV); during animal movement with large astrocyte Ca2+ activity (*violet*: 6.5±2.5 dilations/FOV). All conditions in N = 20 FOV in 6 mice. Compared to dilations during rest, dilations accompanied by large astrocytic Ca2+ activities, alone or simultaneously with movements, were significantly larger (Paired t-test, with Holm correction: Rest vs. Ca2+: p=0.033 and vs Ca2+ during movements: 0.00039. Rest vs movement: p= 0.12. Movement without vs with Ca2+ p=0.062. All values were compared with ANOVA before t-test p=0.0011).

**Sup. Figure S3.**
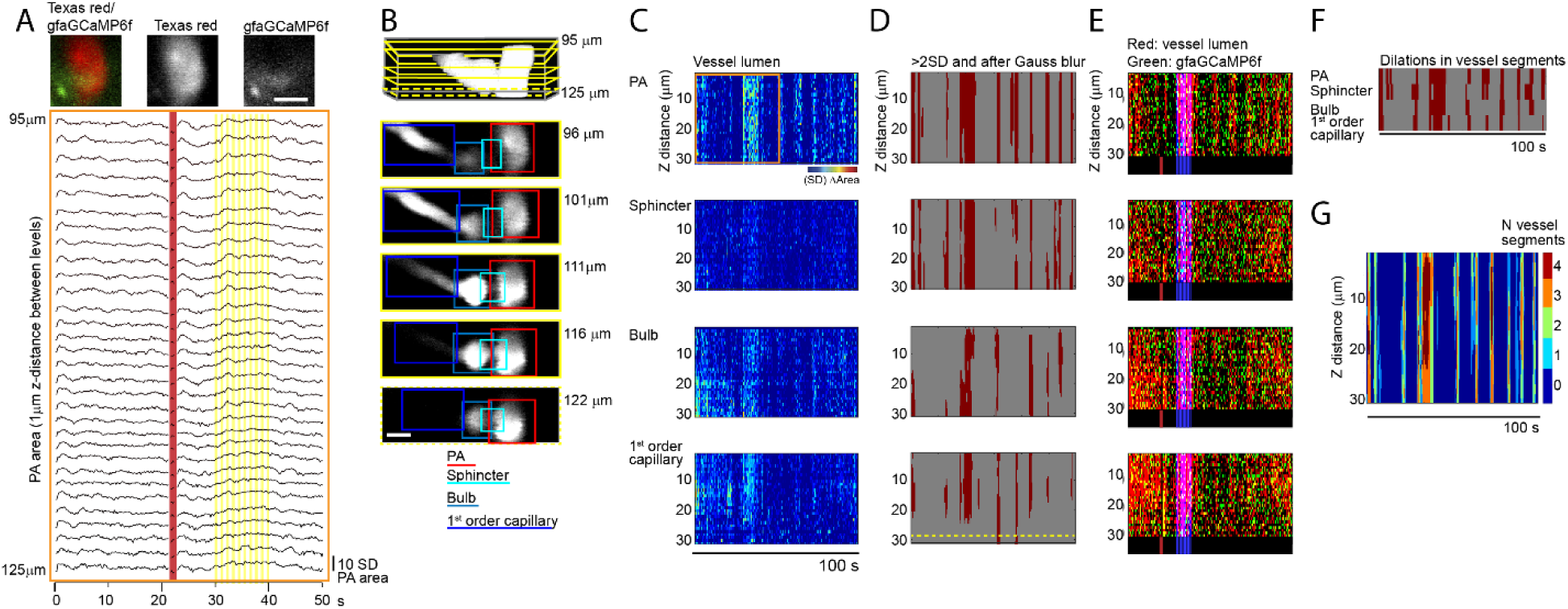
Method to detect dilation patterns in a 3D-FOV. A) *Top*: cumulative average two-photon image of a PA at the capillary branch point and surrounding astrocyte end-feet in a single z-level focal plane in auditory cortex during a tone stimulation experiment. *Red*: Texas red-dextran shows vessel lumen. *Green*: GFAPGCaMP6f highlights astrocyte structures via the cumulative Ca2+ signal during the acquisition. *Left*: color overlay of red and green channels; *middle*: red channel (represented in black and white); *right*: green channel (in black and white). Scalebar: 10 µm. *Bottom:* changes in the PA vessel area shown for all the focal planes recorded in z between 95-125 µm below the surface at 1 µm interdistance. *Red line*: identifies the timing of a dilation throughout the FOV related to a short animal movement; *yellow highlight*: identifies the period of a longer dilation evoked by a tone stimulation. B) *Top:* cumulative average of the entire 3D stack of Texas red-dextran vessel lumen images (*white*) used in panel A and in the rest of this figure. *Yellow rectangles* outline the position in the stack of the 5 z-levels used as examples below. One of the levels is *stippled* as it represents the example also shown in panel D. *Bottom*: rectangular crops of the images at each of the 5 selected z-levels are manually defined as explained in Suppl. Fig. 1C and color-coded to mark the various vessel segments (*red*: PA; *cyan*: sphincter; *light blue*: bulb; *blue*: 1^st^ order capillary). Imaging depths below cortical surface are indicated next to the pictures. Scalebar: 10 µm. C) Representative z-t depiction of the changes in vessel area during a 100 s 3D imaging session for (*top to bottom*) PA, sphincter, bulb and 1^st^ order capillary. The normalized values of changed vessel area measured in the rectangular x-y image crops at each z level are shown in a map of pixel intensity (*jet colormap*) over time. The *orange box* in the PA depiction marks the same period and z-levels presented as traces in panel A. D) Binarization of the data shown in C: periods of dilations (*red*) correspond to periods in which the z-t pixels have >2SD values after Gaussian filter blurring. Notice that the capillary in this example is only present in the top and bottom z-levels during the largest expansions as illustrated by the yellow dotted line at the bottom of the panel. E) Representative z-t depiction of the changes in vessel area during the 100 s 3D imaging session overlaying the normalized vessel area detected with Texas red for each z-level (*red*) and the average GFAP-GCaMP6f Ca2+ signal from astrocyte end-feet surrounding the vessel at the corresponding z-level (*green*). In overlay is also shown in *blue* the period with tone stimulation, giving an overall pinkish appearance. For better visual clarity, the *blue* period is also shown in a black bar below the z-t depiction, together with a *red line* indicating the time of a mouse movement. F) Summary of the dilation periods (*red*) for each of the four vessel compartments shown in D, here presented in a binary form that just displays dilation or non-dilation of the whole compartment. This representation visualizes the timing of all the dilations and highlights that different dilations initiate at different vessel segments. However, it disregards the individual event’s characteristics that are instead presented in D (x-y area of dilation, shown as pixels’ intensity) and in E (z-levels involved in the dilation). G) Overlay of the dilation patterns of the different vessel segments summarized in F. In this case, a color code is used to show the number of vessel segments participating in each dilation at any specific time during the 100 s acquisition period. The graphical representation helps visualizing that not all vessel segments participate in each dilation.

**Sup. Figure S4.**
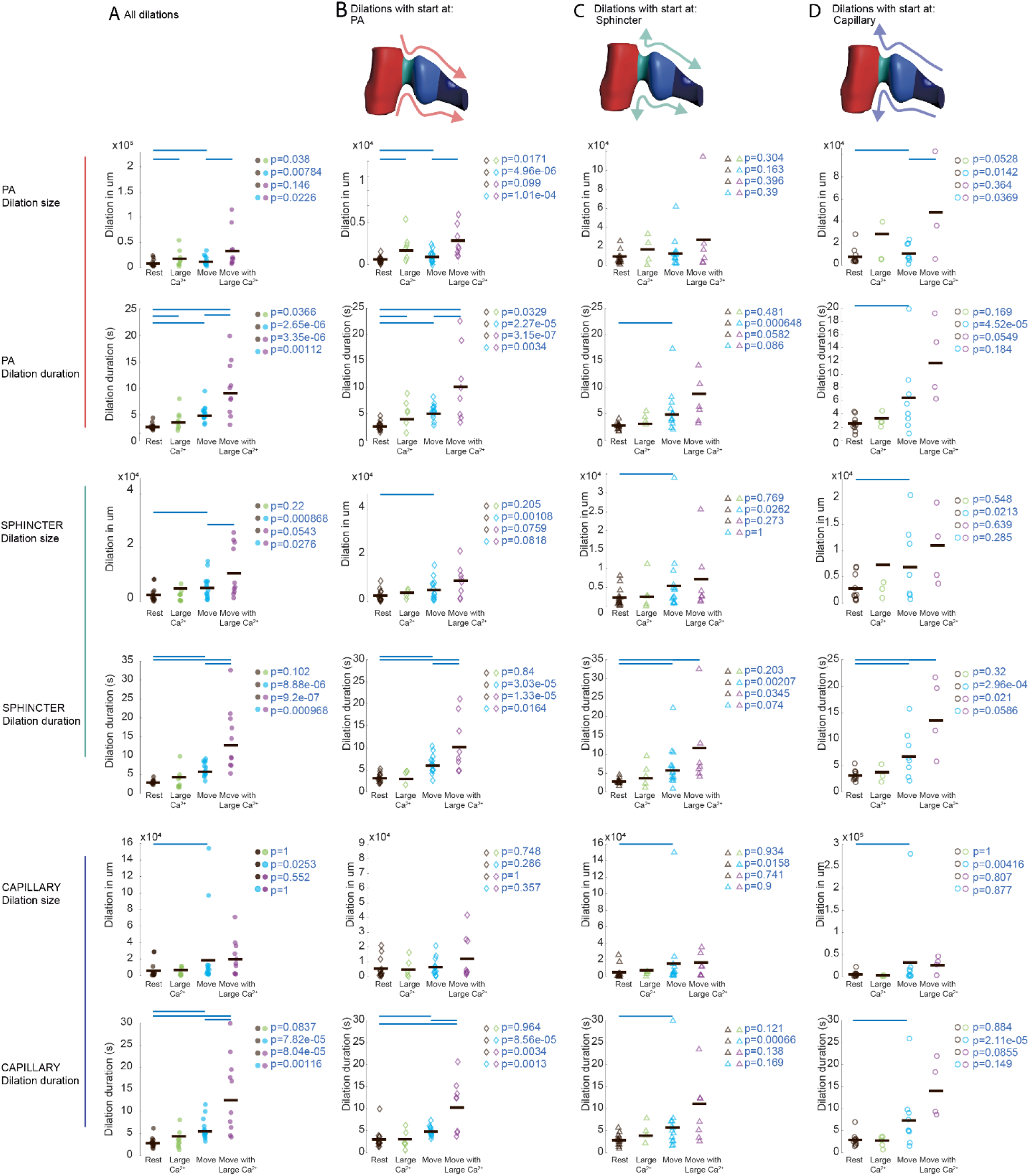
Size and duration of dilations in different vessel segments do not depend on the initiation site but are differently affected by the mouse movements and by large astrocytic Ca2+ activity. A) Column presenting quantitative analysis of the size and duration of all the dilations seen in the different vessel segments (from *top* to *bottom*: PA, sphincter, and capillary) independent of the initiation site but according to the condition in which they occurred, namely during: resting periods of the mouse (*black circles*); resting periods showing large astrocytic Ca2+ activities (*green circles*); periods of movement of the mouse (*azure circles*); periods of movement showing large astrocyte Ca2+ activities (*violet circles*). The main observations are: in the resting mouse, when astrocyte Ca2+ activity was present, size and duration of the dilations increased at the level of the PA, but not of the sphincter and 1^st^ order capillary. When the mouse moved, but no large astrocyte Ca2+ activity was seen, duration and size of the dilations increased in all three segments. The co-occurrence of movement and large astrocyte Ca2+ activity led to an increase in duration of dilations in all vessel segments, but only increased size of dilations in PA and sphincter. Number of dilations/FOV were: during rest: 162±31.2; during rest + large astrocyte Ca2+: 4.2±2.0; during movement without astro Ca2+: 47.6±22.1; during movement + large astrocyte Ca2+: 6.5±2.5. Statistical comparisons were performed with the t-test, modified with Holm correction for multiple comparisons. All comparisons (colored circles) and corresponding p-values are shown in the figure; if p>0.05, a blue line is shown above the two compared datasets. Data from: N=20 FOV in 6 mice. B) Column presenting the same type of analysis as in A but restricted to dilations with a perceived initiation at the PA, i.e. arriving from the PA compartment (symbols: *diamonds*). The results are largely similar to those obtained in A, where we considered all dilations independent of their initiation site. Number of dilations/FOV were: during rest: 50.9±12.8; during rest + large astrocyte Ca2+: 1.92±0.8; during movement without astro Ca2+: 15.0±7.1; during movement + large astrocyte Ca2+: 2.1±0.9. Statistical comparisons were performed and are presented as in A. C) Column presenting the same type of analysis as in A and B but here restricted to dilations starting at the sphincter (symbols: *triangles*). These dilations do not apparently show sensitivity to large astrocytic Ca2+ activities but are affected by movements similarly to what seen in A. Number of dilations/FOV were: during rest: 40.3±8.5; during rest + large astrocyte Ca2+: 1.6±0.5; during movement without astro Ca2+: 12.4±6.9; during movement + large astrocyte Ca2+: 2.3±1.0. Statistical comparisons were performed and are presented as in A. D) Column presenting the same type of analysis as in A, B and C, but here considering just dilations with a perceived initiation at the capillary level, i.e. arriving from the capillary compartment (symbols: *circles*). These dilations, like those starting at the sphincter, do not appear to be strongly affected by large astrocytic Ca2+ activity, but are affected by movements similarly to what seen in A. Number of dilations/FOV were: during rest: 56.4±13.0; during rest + large astrocyte Ca2+: 2.1±0.8; during movement without astro Ca2+: 14.7±7.9; during movement + large astrocyte Ca2+: 2.4±0.7. Statistical comparisons were performed and are presented as in A.

**Sup. Figure S5.**
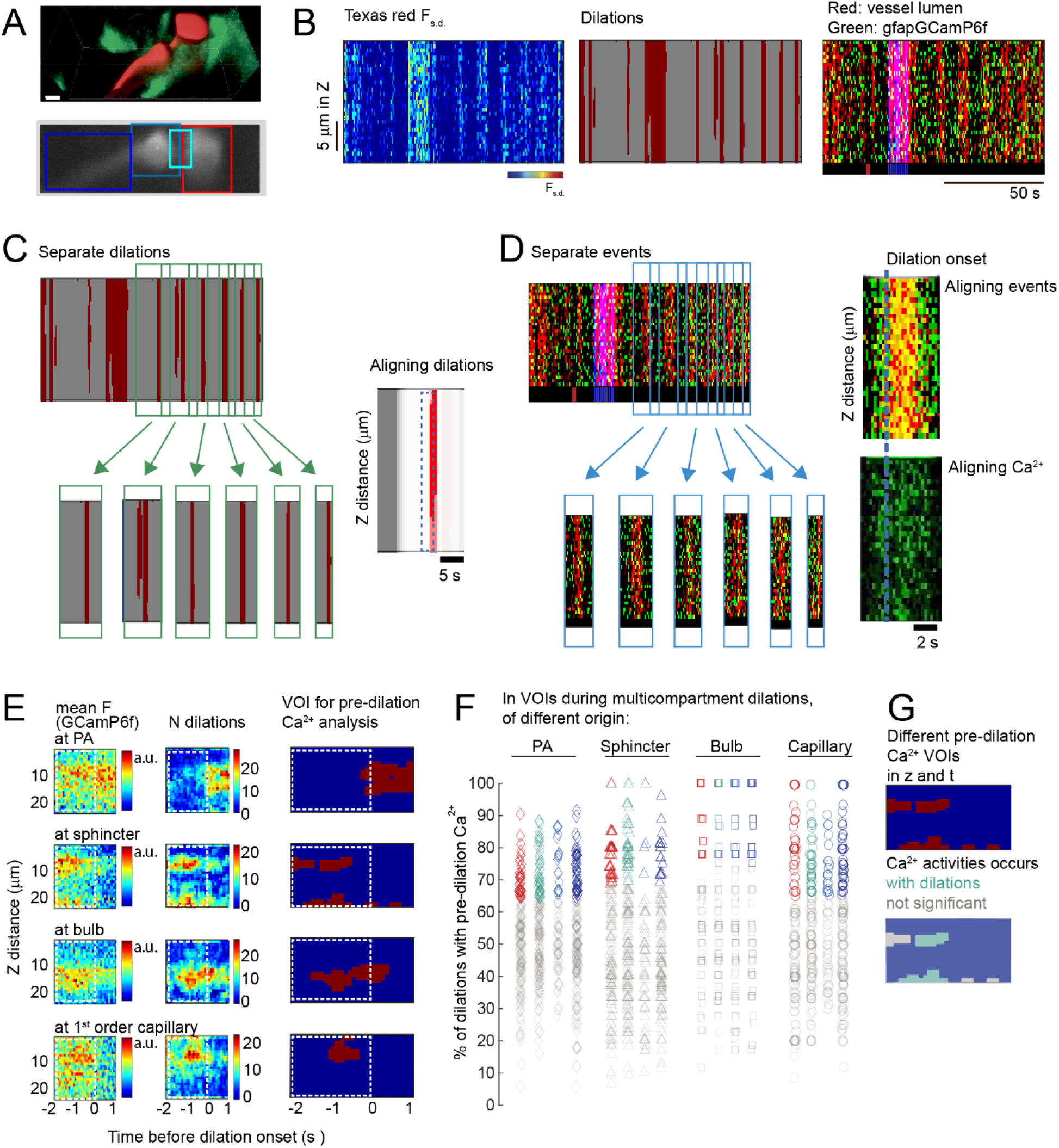
Method to detect regions with pre-dilation astrocytic Ca2+ activity. A) Cumulative average of a 3D timeseries from the auditory cortex of a Texas red-dextran-injected *GFAPCreERT2:GCaMP6f* mouse, revealing a 1^st^ order capillary branch from a PA and neighboring astrocyte structures. *Top*: entire 3D stack taken 64-86 µm below the cortical surface. *Red*: masked vessel lumen. *Green*: GFAPGCaMP6f signal outlining the surrounding astrocyte structures. *Bottom*: an individual focal plane from the same stack, taken 72 µm below the surface. The average Texas red-dextran signal from the vessel lumen in black and white and four manually drawn color-coded rectangles identify the different vessel segments: *red*: PA; *cyan*: sphincter; *light blue*: bulb; *blue*: capillary. Scalebar: 5µm. B) Representative z-t depiction of the dilation events of the PA during the 100 s 3D imaging session across the different z-levels. Data are presented as in Fig S3C-E (see there for more details). *Left*: normalized values of changed vessel area measured in the rectangular x-y image crops at each z level are shown in a map of pixels intensity (*jet colormap*) over time. *Middle*: the data on the left are binarized to depict the periods of dilation (*red*) and the level of expansion of the dilations along the z-axis. *Right*: normalized signal from Texas red (*red*) and GFAPGCaMP6f (*green*). *Blue*: period indicates tone stimulation. *Red* in the black bar below the z-t depiction identifies a movement of the mouse. C) *Left*: Periods with dilation events detected in the z-t rendering were considered separate events (see criteria in Methods, section “*Detection of dilation events and patterns”*) and cropped (*green arrows* and *boxes*). *Right*: as in the example, all the crops where temporally aligned based on the onset time of each dilation to allow for detection of recurrent “pre-dilation” astrocyte Ca2+ activity in a 3-sec period including the 2-sec preceding dilation onset (*blue dashed line box)* and the first second after dilation start, the latter to avoid abrupt cutting Ca2+ dynamics initiated within the 2-sec pre-dilation period. D) Same data as in B, showing on the *left* the normalized vascular signal from Texas red (*red*) and astrocyte end-foot Ca2+ from GFAPGCaMP6f (*green*). Data related to separate dilations are cropped and temporally aligned as in C. *Right*: expanded example of the average Texas red and GFAP-GCaMP6 signals (*top*), or only the latter (*bottom*), showing that astrocytic Ca^2+^ activity rises before start of dilation (*dotted blue line*) and lasts through the dilation. E) Defining the z-t position of astrocytic pre-dilation Ca^2+^ activity in different end-feet ROIs on the vessel segments. The 2-sec period before dilation onset is marked with a white dashed box. *Left*: mean GFAP-GCaMP6f signal (*pseudocolor scale*) in astrocyte end-feet during the relevant 3-sec period in relation to vessel segments. *Middle*: the *pseudocolor scale* indicates in this case the number of dilations (N) during all imaging sessions showing an astrocyte Ca2+ elevation at this time and depth in z. The number of Ca2+ elevations was defined by summing up all the individual masks created from the mean GFAPGCaMP6f signals. *Right*: Binarization of the Ca2+ signals after Gaussian filtering; the resulting mask (*red* in *blue* background) defines the putative recurrent “pre-dilation Ca^2+^ signals” as voxels of interest (VOI) in relation to cortical depth and time relative to dilation onset. This mask was then used to evaluate for each individual dilation the presence or not of these specific VOIs of pre-dilation Ca2+ activity, as described in methods section” *Detection and analysis of pre-dilation astrocytic 3D Ca2+ activity*”. F) Percentage of dilations preceded by pre-dilation end-foot Ca2+ activity in VOIs located at the level of different vessel segments, according to the segment of origination of the dilation. The colors of the data points indicate the vessel segment at which the astrocytic end-feet VOI is located (*red*: PA; *cyan*: sphincter; *light blue*: bulb; *blue*: capillary). The symbols indicate the vessel segment of origin of the dilation (*diamond*: PA; *triangle*: sphincter; *square*: bulb; *circle*: capillary). The level of prevalence was used to define whether pre-dilation Ca2+ activities could be considered concurrently present in relation to a given dilation pattern. They were considered present if they occurred with 100 % of dilations subtracted of the 0.1 confidence limit, which depended on the number of dilations of a given dilation pattern (see Methods). The *colored* data points mark the pre-dilation Ca2+ activities that were considered statistically present before dilations of a specific pattern. The grey data points mark the activities that were not counted. G) Illustration of the statistical attribution of a group of astrocyte Ca2+ events to the specific sub-group of pre-dilations Ca^2+^ events. *Top*: binarized astrocyte Ca2+ elevation VOIs at the sphincter shown in E. *Bottom*: the Ca2+ events occurring consistently before all dilations (*cyan* because around sphincter) were assigned to the “pre-dilation” sub-group (see F), the others appearing randomly before only some of the dilations were not included in the “pre-dilation” events (*grey*).

**Sup. Figure S6.**
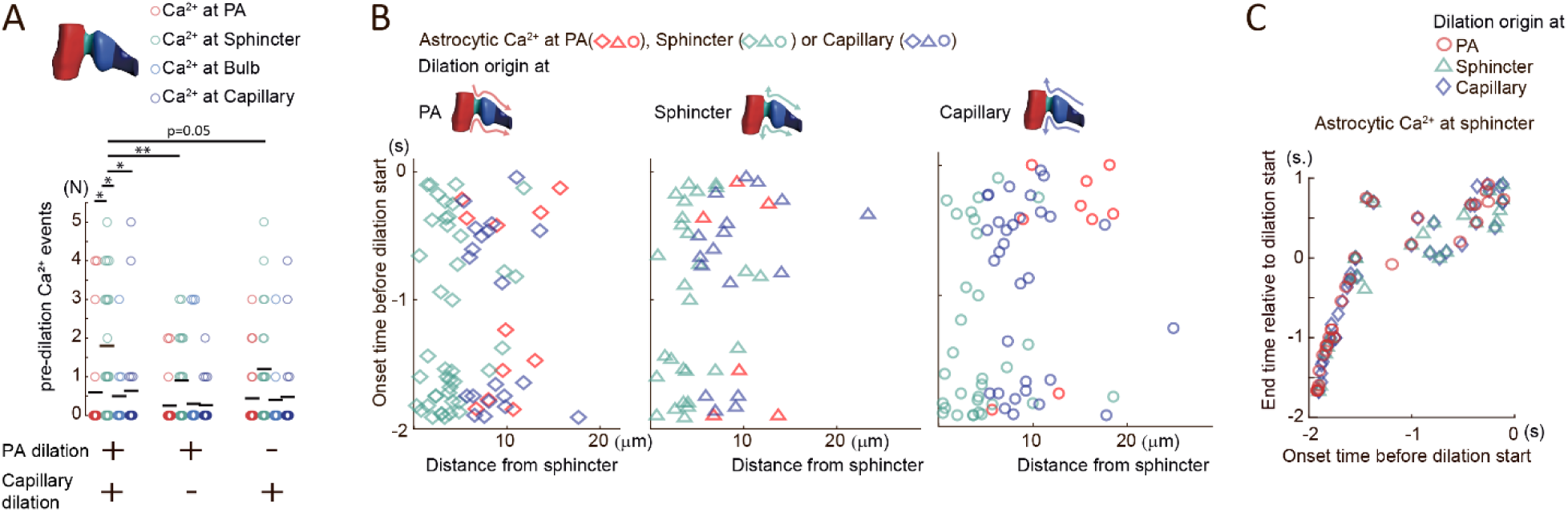
Pre-dilation astrocytic end-foot Ca^2+^ activity consisting of small local, not-spreading events occurs earlier and more frequently at the sphincter during multicompartmental dilations. A. Pre-dilation astrocytic end-foot Ca2+ activity during different types of dilations, either multi-compartmental, or mono-compartmental involving just the PA, or the 1^st^ order capillary branch. The number of pre-dilation Ca2+ events occurring at the sphincter is significantly higher than those occurring at all the other compartments during multicompartmental dilations (*left*) but not significantly so during dilations of just the PA (*middle*) or the capillary branch (*right).* The number of events at the sphincter is also higher in case of multicompartmental dilations vs monocompartment dilations, independent if the latter involve the PA or the 1^st^ order capillary branch. Statistical analysis: Kruskal Wallis ANOVA followed by Wilcoxon test with Holm-correction for multiple comparisons: (a) Multicompartmental dilations: number of end-foot Ca2+ events at sphincter (*cyan circles*) vs. at PA (*red*): p=0.012; at sphincter vs. at bulb (*light blue*): p=0.017; at sphincter vs. at capillary (*blue*): p=0.0178. After Kruskal Wallis test, the number of Ca2+ events at sphincter in multicompartmental dilations is significantly different from the number in monocompartment dilations involving just the PA (p=0.0096), or in monocompartment dilations involving just the 1^st^ order capillary branch (p=0.05). Data are from 82.7±20.1 multicompartmental dilations/FOV, 15.3±4.5 monocompartment dilations involving just the PA/FOV; and 61.4±14.7 mono-compartment dilations involving just the capillary branch/FOV; N=20 FOV in 6 mice. B. Onset time of pre-dilation astrocytic Ca2+ activity according to distance from the sphincter. Localization of the astrocyte end-foot Ca2+ activity: at PA (*red*); at sphincter (*cyan*); at capillary (*blue*). Origin of the dilation: at PA (*diamonds*); at sphincter (*triangles*); at capillaries (*circles*). The distribution shows that the astrocyte Ca2+ activity preceding these multicompartment dilations does not spread from the sphincter like a wave. Experimental points are from 116.1±24.7, 82.4±19.7 and 67.2±16.2 dilations/FOV for dilations starting at PA, sphincter and capillary, respectively. From N=20 FOVs in 6 mice. C. Correlation between onset-time and end-time of pre-dilation Ca2+ activities at the sphincter before multicompartmental dilations arriving from either the PA (*red circles*) or the capillary (*blue diamonds*) or initiated at the sphincter (*cyan triangles*). The graph shows two groups of time-correlated activities, fully preceding or persisting after dilation onset.

**Sup. Figure S7.**
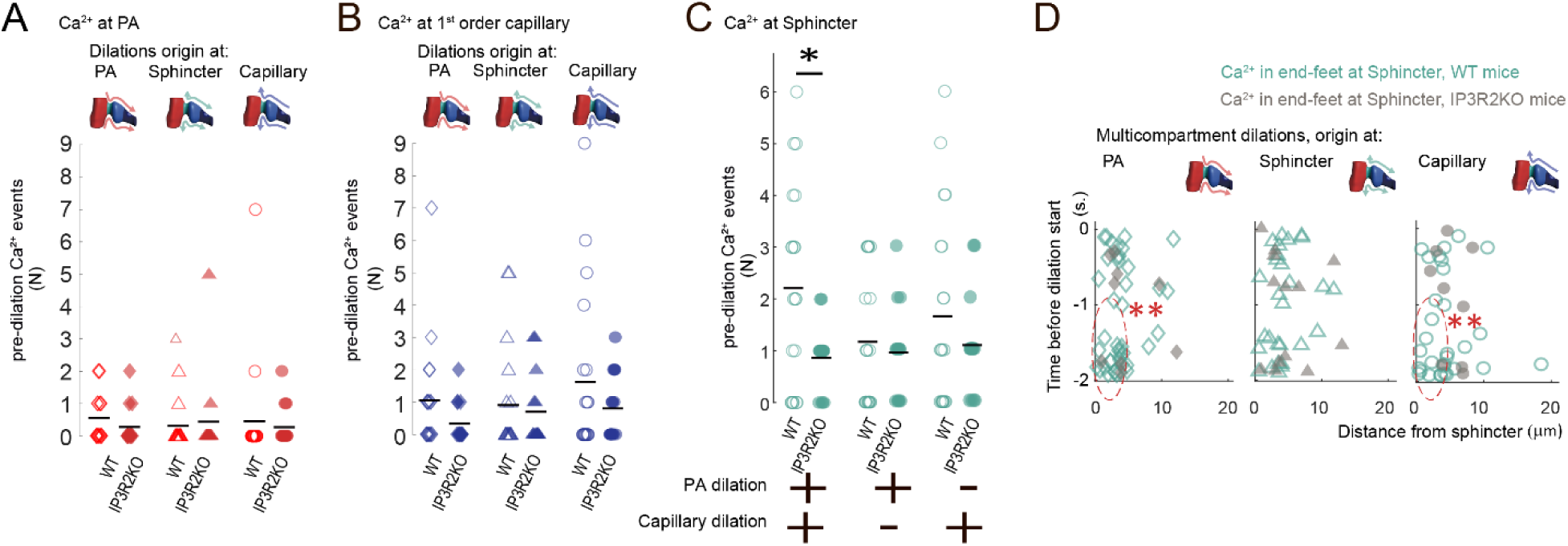
IP3R2-dependent and IP3R2-independent pre-dilation astrocytic end-foot Ca^2+^ activity depending on type of dilation and location of the active end-feet. A. Number of pre-dilation astrocytic end-foot Ca2+ events seen at the PA in WT (*red, open symbols*) vs. IP3R2KO (*red, solid symbols*) mice in multicompartmental dilations arriving from the PA (*diamonds*), or the capillaries (*circles*), or initiated at the sphincter (*triangles*). No statistical difference was found in the number of Ca2+ events in WT vs. IP3R2KO mice in all three the dilation patterns (Mann-Whitney test). Specifically: dilations originating at PA: p=0.11; WT: 116.1±24.7 dilations/FOV; IP3R2KO: 130.3±35.4 dilations/FOV. Dilations originating at the sphincter: p=0.89; WT: 82.4±19.7 dilations/FOV; IP3R2KO: 76.9±21.9 dilations/FOV. Dilations originating at capillary: p=0.55; WT: 67.2±16.2 dilations/FOV; IP3R2KO: 50.9±14.3 dilations/FOV. Data in WT: N=20 FOVs in 6 mice; in IP3R2KO: N=14 FOV in 4 mice. B. Number of pre-dilation astrocytic end-foot Ca2+ events seen at the capillaries in WT (*blue, open symbols*) vs. IP3R2KO (*blue, solid symbols*) mice in multicompartment dilations arriving from the PA (*diamonds*) or the capillaries (*triangles*) or initiated at the sphincter (*circles*). No statistical difference was found in the number of Ca2+ events in WT vs IP3R2KO mice in all three the dilation patterns (Mann-Whitney test). Specifically: dilations originating at PA: p=0.055; WT: 116.1±24.7 dilations/FOV; IP3R2KO: 130.3±35.4 dilations/FOV. Dilations originating at the sphincter: p=0.44; WT: 82.4±19.7 dilations/FOV; IP3R2KO: 76.9±21.9 dilations/FOV. Dilations originating at capillary: p=0.35; WT: 67.2±16.2 dilations/FOV; IP3R2KO: 50.9±14.3 dilations/FOV. Data in WT: N=20 FOVs in 6 mice; in IP3R2KO: N=14 FOV in 4 mice. C. Number of pre-dilation astrocytic end-foot Ca2+ events at the sphincter in WT (*cyan, open circles*) vs. IP3R2KO (*cyan, solid circles*) mice in multicompartmental dilations, and in monocompartment dilations either involving just the PA or the capillary. This type of information concerning the monocompartmental dilations was not presented in Figure 4C and is added here for comparison with the IP3R2KO effects on multicompartmental dilations. During the latter, the number of pre-dilation Ca2+ events in end-feet around the sphincter was significantly lower in IP3R2KO vs. WT mice (Mann-Whitney test: p=0.028; WT: 82.7±20.1 dilations/FOV; IP3R2KO: 75.3±16.7/FOV). In contrast, in dilations that did not spread across the sphincter, the number of pre-dilation Ca2+ events was not significantly different in IP3R2KO and WT mice (Mann-Whitney test). Specifically: dilations involving just the 1^st^ order capillary branch: p=0.36. WT: 61.4±14.7 dilations/FOV; IP3R2KO: 83.3±25.6 dilations/FOV. Dilations involving just the PA : p=0.8. WT: 15.3±4.5 dilations/FOV; IP3R2KO: 10.3±2.5 dilations/FOV. Data in WT: N=20 FOVs in 6 mice; in IP3R2KO: N=14 FOV in 4 mice. D. Comparison of the onset time and distance from the sphincter of the astrocyte pre-dilation Ca2+ events for multicompartmental dilations in IP3R2KO (*grey, solid symbols*) vs WT (*cyan, open symbols*) mice. The two parameters are compared according to the origin of the dilations to which they are associated: at either the PA (*diamonds*), the capillary (*circles*), or the sphincter (*triangles*). The pre-dilation Ca2+ signals suppressed in IP3R2KO mice were mostly those occurring early and closest to the sphincter. As reported in the legend to Figure 4, for both dilations starting at PA and capillary, the Ca2+ events occurring at short distance from the sphincter (SD, <5 µm) were significantly more reduced in IP3R2KO vs WT (*dashed red circles*) than those occurring at long distance (LD, 5-20 µm). Statistics: Mann-Whitney test: WT vs KO: for dilations starting: (a) at PA: SD: p=0.0038; LD: p=0.18; (b) at sphincter: SD: p=0.52; LD: p=0.18; (c) at capillary: SD: p=0.023, LD: p=0.15. The number of dilations with origin at PA, sphincter and capillary were, respectively: in WT: 116.1±24.7, 82.4±19.7 and 67.2±16.2 in 20 FOVs in 6 mice; in IP3R2KO: 130.3±35.4, 76.9±21.9 and 50.9±14.3 in 14 FOVs in 4 mice.

**Sup. Figure S8.**
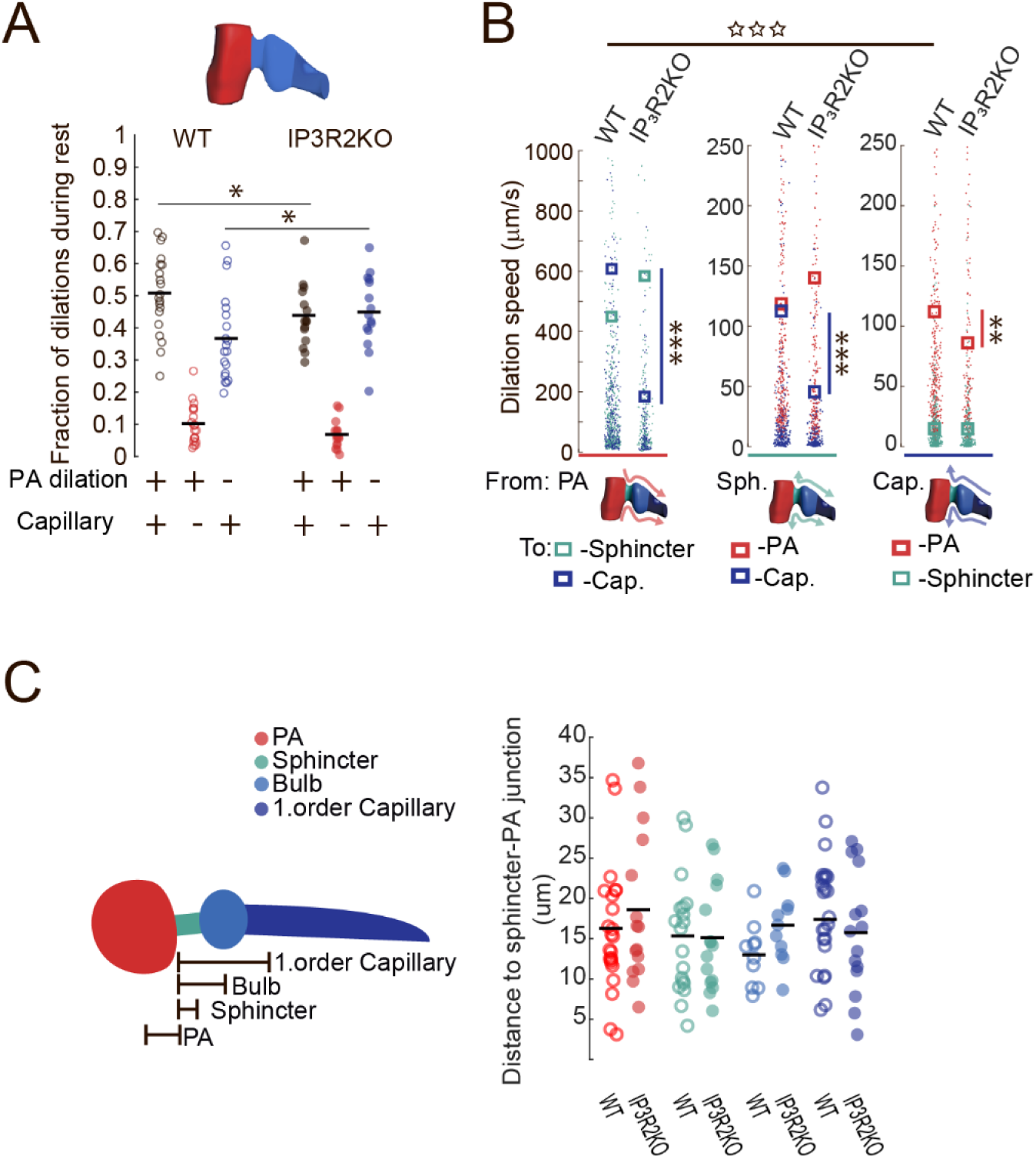
Dilation patterns and speed of dilation spread in IP3R2KO vs WT mice. A) Fraction of multicompartmental (*black*) and monocompartmental (either of PA, *red*, or capillary, *blue*) dilations in WT (*open circles*) vs IP3R2KO (*solid circles*) mice at rest. In IP3R2KO mice, multicompartmental dilations significantly decreased compared to WT mice, whereas dilations involving just the capillaries increased. Statistical analysis: Friedman ANOVA for each group followed by X^2^ test comparing WT against IP3R2KO and Mann-Whitney test: multicompartmental dilations: WT vs IP3R2KO: p=0.037; monocompartmental dilations involving the 1^st^ capillary branch: WT vs IP3R2KO: p=0.02; WT: 236±45 dilations/FOV; N=20 FOV, 6 mice; IP3R2KO: 235±53 dilations/FOV; N=14 FOV; 4 mice. B) Speed by which dilations spread between vessel compartments in WT and IP3R2KO mice according to the site of origination of the dilation (*left*: PA, *red*; *middle*: sphincter, *cyan*; *right*: capillary, *blue*). Color-coded dots along the y-axis mark the dilation spread time of a given vessel compartment, measured as the distance from the initiating compartment in µm/the delay from the start time. The color-coded rectangles are the corresponding mean values (*red*: at PA; *cyan*: at sphincter; *light blue*: at bulb; *blue*: at capillary). *Rectangles* indicate the mean speed value for spread in a given compartment, color-coded as above. In WT mice, dilations spread at much faster speed between PA and capillary compartments than vice versa (open stars on the top): Mann-Whitney test: PA to capillary (*blue points* in plot to the *left*) vs capillary to PA (*red points* in plot to the *right*): p=1.9x10^-5^; PA to capillary: N=366 dilations; capillary to PA: N=323 dilations; 20 FOVs; 6 mice). In IP3R2KO mice, dilations between PA and capillary spread at significantly reduced speed than in WT mice, regardless of whether: (a, *left*) the dilation arrived from the PA (Mann-Whitney test, WT vs IP3R2KO: dilation from PA to sphincter: p=0.15; or to capillary: p=1.5x10^-7^; WT: N=366 dilations, IP3R2KO: N=142 dilations); (b, *middle*) the dilation was initiated at the sphincter (Mann-Whitney test: WT vs IP3R2KO: dilation from sphincter to PA p=0.79 or to capillary p=6.02 x 10^-7^; WT: N=329 dilations; IP3R2KO: N=239 dilations); or (c, *right*) the dilation arrived from the capillary bed (Mann-Whitney test: WT vs IP3R2KO: dilation from capillary to sphincter p=0.96; and to PA p=0.0076; WT: N=323 dilations; IP3R2KO: N=145 dilations). Data for WT from N=20 FOV in 6 WT mice, and for IP3R2KO form N=14 FOV in 4 mice. C) Comparison of the average distance between vessel compartments in WT and IP3R2KO mice. *Left*: schematic representation of how the distance between each vessel compartment and the connection point between PA and sphincter was calculated for each FOV included in the study. The distances were relative to the point in which the sphincter met the PA circumference using as reference positions the center of the circular PA and bulb structures, and the middle point of the rectangular sphincter and capillary structures. *Red*: PA; *cyan*: sphincter; *light blue*: bulb; *blue*: capillary. *Right*: comparison of the distance between each vessel compartment and the connection point between PA and sphincter in WT (*open circles*) and IP3R2KO (*solid circles*) mice. Compartments identified by same colors as in *left*. No difference was found in WT vs IP3R2KO mice, though a large diversity was observed in the measures in each group. Paired t-test WT vs IP3R2KO: distance from the junction between PA and sphincter to the center of the PA: p=0.42; to the middle of the sphincter segment: p=0.92; to the center of the bulb: p=0.068; to the middle of the capillary segment: p=0.52 Data from N=20 FOV, 6 WT mice, N=14 FOV, 4 IP3R2KO mice.

**Sup. Figure S9.**
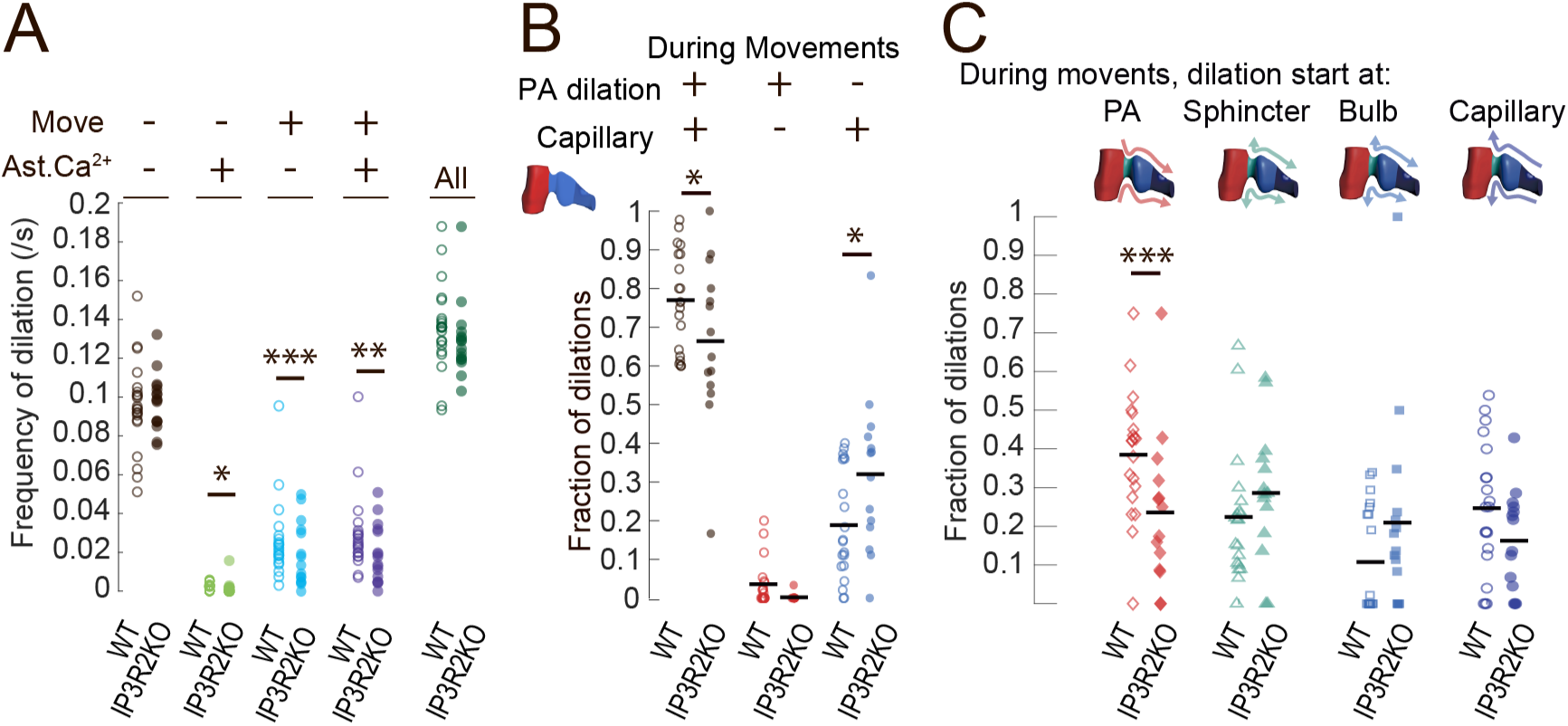
Comparative analysis of dilation features in different conditions in IP3R2KO vs WT mice. A) Frequency of all dilations independent of origin, direction and pattern, analyzed in different conditions in WT (*open circles*) and IP3R2KO (*solid circles*) mice. Conditions are: mouse at rest (*black*); mouse at rest displaying large astrocyte Ca2+ activity (*green*); mouse showing movements (*azure*); mouse showing movements and large astrocyte Ca2+ activity (*violet*). Moreover, to the far right, are the frequency values summing up all the dilation events in all conditions (*dark green*). Across all conditions, a X^2^-test did not indicate difference in dilation frequencies (p=0.39). Moreover, no frequency difference appeared between WT and IP3R2KO mice with the Mann-Whitney test when all the dilation events were compared independently of whether the mouse was moving, resting etc: WT vs IP3R2KO: p=0.16. In contrast, comparison of the dilations taking into account the specific condition, revealed that frequency of dilation was reduced in IP3R2KO for dilations that occurred during animal movements or were accompanied by large astrocyte Ca2+ activities or by both, but not for dilations in the resting animal. Mann-Whitney test: WT vs IP3R2KO: at rest: p=0.85; at rest + large Ca2+ activity: p=0.036; moving mouse: p=6.2x10-7: moving mouse + large Ca2+ activity: p=0.0059. In WT mice, the dilations/FOV/condition were: at rest: 162±31.2; at rest + large Ca2+ activity: 4.2±2.0; moving mouse: 47.6±22.1; moving mouse + large Ca2+ activity: 6.5±2.5; N = 20 FOV in 6 mice. In IP3R2KO mice the dilations/FOV/condition were: at rest: 176±39.8; at rest + large Ca2+ activity: 2.27±1.61; moving mouse: 34.1±15.2; moving mouse + large Ca2+ activity: 1.73±1.28; N = 14 FOV in 4 mice. B) Fraction of dilations involving different vessel compartments in WT (*open circles*) vs IP3R2KO (*solid circles*) mice during mouse movements. Multicompartmental dilations (*black circles*) were reduced, while monocompartmental ones including just capillaries (*blue circles*) were increased in IP3R2KO vs WT mice during movements (Friedman ANOVA followed by X^2^ test comparing WT vs IP3R2KO and Mann-Whitney test: multicompartmental: WT vs IP3R2KO: p=0.045; just capillary: WT vs IP3R2KO: p=0.017). The fraction of monocompartment dilations involving just the PA (*red circles*) did not significantly change in IP3R2KO vs WT. In WT mice during movement: 47.6± 22.1 dilations/FOV; N=20 FOV, 6 mice; In IP3R2KO mice during movement: 34.1± 15.2 dilations/FOV N=14 FOV, 4 mice. C) Fraction of multicompartmental dilations according to origination site seen during mouse movements in WT (*open symbols*) and IP3R2KO (*solid symbols*) mice. *Red diamonds*: dilations originating at PA; *cyan triangles*: at sphincter; *light blue squares*: at bulb; *blue circles*: at capillary. Changes in dilation patterns mirror those found in the resting mouse (Figure 5A), but here only the fraction of multicompartmental dilations beginning at the PA was significantly reduced in the IP3R2KO mice. (Friedmans ANOVA, followed by Mann-Whitney test: WT vs IP3R2KO: start at PA: p=0.0041; at sphincter: p=0.059; at bulb: p=0.13; at capillary: p=0.070; In WT mice during movement: 35.7± 14.08 dilations/FOV N=20 FOV in 6 mice; In IP3R2KO mice during movement: 23.14± 8.86 dilations/FOV N=14 FOV in 4 mice.

**Sup. Figure S10.**
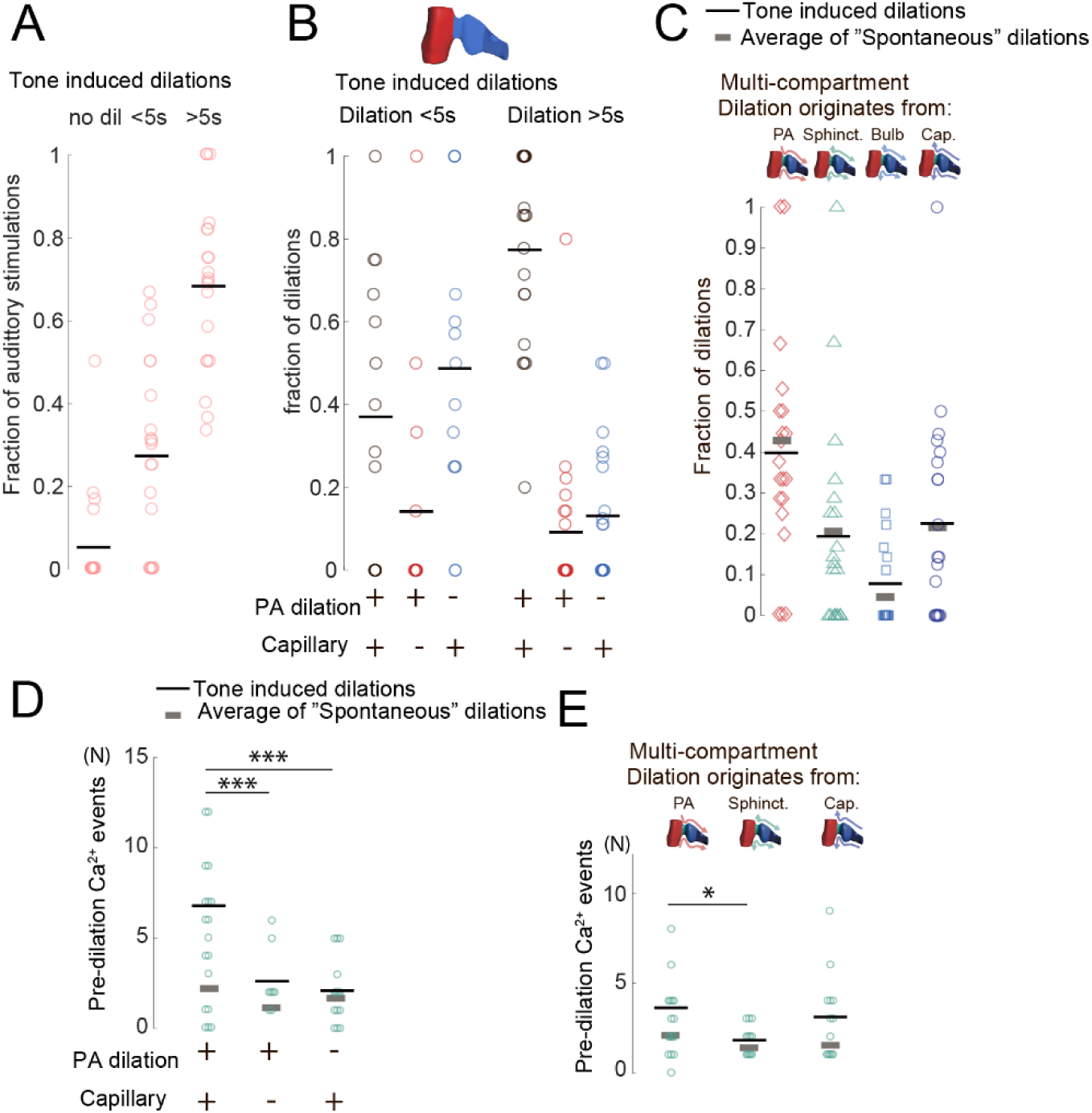
Dilation patterns in auditory cortex were similar for tone-evoked and spontaneous dilations. A) Temporal distribution of dilation responses to our protocol of auditory stimulations expressed as a fraction of all the auditory stimulations. Only a few stimulations did not trigger any dilation response (no dil). On average, ∼ 30% of the stimulations triggered dilations <5s, and >65% triggered dilations >5s. Data are: 7.55±1.79 dilations/FOV; N=20 FOV; 6 mice. B) Fraction of tone-evoked dilations that are multicompartmental (*black circles*), or monocompartmental, the latter sub-divided in those involving the PA (*red circles)* or the 1^st^ order capillary (*blue circles*). The fractions are presented in two groups, separated based on their duration, either <5S or >5s and compared between these groups. As seen also with spontaneous dilations, the proportion of multicompartment dilations induced by tone stimulations was higher among longer-lasting dilations: <5s dilations were more often restricted to just the capillary bed, whereas >5s dilations most often included all vessel compartments. Friedmans ANOVA for each response duration followed by comparison using the X^2^ test. Dilations <5s: p=0.062, X^2^=5.6; dilations >5s: p=4.7x10^-6^, X^2^=24.5. ANOVA shows unequal distribution of dilations only in the >5s dilations group. The X^2^ values support this difference between shorter and longer dilations. Data are: 2.15±0.92 <5s dilations/FOV and 5.4±1.33 >5s dilations/FOV; N=20 FOV in 6 mice. C) Fraction of tone-evoked multicompartmental dilations according to their origination site: at the PA (*red diamonds*), at the sphincter (*cyan triangles*), at the bulb (*light blue squares*), or at the capillary (*dark blue circles*). The average values of tone-induced dilations in each group (*black lines*) are plotted for comparison together with the corresponding average values (*grey lines*) of spontaneous dilations taken from the *ad hoc* set of experiments. Tone-evoked and spontaneous multicompartmental dilations displayed similar proportions of dilations with the same origination. Friedmans ANOVA. followed by Wilcoxon test: spontaneous vs tone-evoke: originating at PA: p=0.49; at sphincter: p=0.25; at bulb: p=0.68; at capillary: p=0.67. The number of spontaneous dilations/FOV analyzed was 82.65±20.13 and 4.15±1.13 tone induced dilations/FOV was in N=20 FOVs in 6 mice. D) Number of FOVs showing pre-dilation astrocytic end-foot Ca2+ activity at the sphincter in relation to tone-evoked dilation patterns, either multi-compartmental or involving just the PA or the capillary branch. The average values for tone-evoked dilations (*black lines*) are plotted for comparison together with the corresponding average values from spontaneous dilations taken from the *ad hoc* set of experiments (*grey lines,* see also supplemental Figure S6A). Tested with Kruskal Wallis ANOVA (p=0.00013) followed by Wilcoxon test. For tone-evoked dilations, we found a significantly higher number of pre-dilation Ca2+ activity at sphincter in multicompartmental dilations vs monocompartmental ones, both vs those involving just the PA (p=0.00069) and vs those involving just the 1^st^ order capillary branch (p=0.00047). When we compared the distributions of pre-dilation Ca2+ activities in spontaneous vs tone-evoked dilations with Kruskal Wallis ANOVA and X^2^ test, we did not find a difference between the two groups. Data are: tone-evoked dilations/FOV: multicompartmental: 0.6±0.46; monocompartmental involving just PA: 0.65±0.36; or just capillary branch: 4.15±1.13. Spontaneous dilations/FOV: multicompartmental: 82.7±20.1; monocompartmental involving just PA: 61.4±14.7; or just capillary branch: 15.3±4.5. N=20 FOV, 6 WT mice. E) Number of FOVs showing pre-dilation astrocytic end-foot Ca2+ activity at the sphincter associated to multicompartmental dilations, according to their site of origination. The average values for tone-evoked dilations (*black lines*) are plotted for comparison together with the corresponding average values from spontaneous dilations taken from the *ad hoc* set of experiments (*grey lines*). Pre-dilation astrocytic Ca2+ activity was numerous for both tone-evoked and spontaneous dilations, and their proportions according to the site of origination of the dilation was similar for tone-evoked and spontaneous dilations (see also supplemental Figure 3B). Tested with Kruskal Wallis ANOVA (p=0.07) followed by Wilcoxon test. Ca2+ at sphincter in multicompartment dilations starting at sphincter vs starting at PA: p=0.015; or vs starting at capillary: p=0.18. Comparison of the distributions of pre-dilation Ca2+ activities in spontaneous and tone-evoked dilations with Kruskal Wallis ANOVA and X^2^ test did not indicate differences between the two groups as the X^2^ values were of comparable size. Specifically: (a) dilations starting at the PA: spontaneous: p=0.019, X^2^=9.99; tone-evoked: p=0.056, X^2^=7.48. (b) dilations starting at the sphincter: spontaneous: p=0.031, X^2^=8.85; tone-evoked: p=0.11, X^2^=6.13; (c) dilations starting at the capillary: spontaneous: p=0.0032, X^2^=13.75; tone-evoked: p=0.027, X^2^=9.15. Data are: tone-evoked dilations/FOV: starting at PA: 1.98±0.62; at sphincter: 0.85±0.38; at bulb: 0.45±0.30, and at capillary: 1.25±0.58. Spontaneous dilations/FOV: starting at PA: 116.1±24.7; at sphincter: 82.4±19.7; at bulb: 70.4±18.9: at capillary: 67.2±16.2. N=20 FOVs in 6 mice.

## Notes

### Competing Interest Statement

The authors have declared no competing interest.

### Summary of Updates

This manuscript has been improved on several issues to correct minor mistakes and improve clarity with regards to description of data analysis and conclusions.

